# Epicardial extracellular vesicles modulate gene expression following ischemia-reperfusion injury in heart-on-a-chip

**DOI:** 10.1101/2025.07.31.667965

**Authors:** Karl T. Wagner, Dawn Bannerman, Qinghua Wu, Simon Pascual-Gil, Ian Fernandes, Luis Felipe Jiménez Vargas, Shuo Yan, Ramak Khosravi, Sargol Okhovatian, Dhana Abdo, Milica Radisic

## Abstract

The epicardium is an essential regulator of cardiac development, homeostasis, and injury, yet the composition and impacts of epicardial cell-secreted extracellular vesicles (EVs) remain incompletely understood. Here, we harness an epicardial Biowire platform integrating human stem cell derived epicardial cells with functional myocardium and apply transcriptomics to reveal enriched EV transport in tissues containing epicardial cells. Profiling epicardial-EVs identified key miRNAs and their conservation through stimulated epithelial-to-mesenchymal transition. Supplementation of epicardial-EVs to tissues undergoing ischemia–reperfusion injury influenced gene expression associated with reduction of extracellular matrix remodeling, fibroblast activation, and suppression of cell-ECM interactions. Correlation of EV-miRNAs with mRNA targets highlighted the role of miR-30d-5p, miR-9-5p, miR-16-5p, and the let-7 family in moderating deleterious fibrotic activation and matrix remodelling during myocardial injury in vitro.

**Teaser:** Cells from the heart’s outer surface secrete tiny packages of biomolecules that influence how the heart responds to injury

## Introduction

The epicardium, the outermost cellular layer of the heart, fulfills diverse and essential roles in cardiac development, adult homeostasis, and responses to injury. During embryogenesis, subsets of epicardial cells undergo an epithelial-to-mesenchymal transition (**EMT**), losing polarity and intercellular adhesion and migrating into the myocardial layer.(*1, 2*) The resulting cells differentiate into key cardiac lineages, including vascular smooth muscle cells (VSMCs) and cardiac fibroblasts (CFs), and secrete paracrine factors that orchestrate myocardial growth, cardiomyocyte (CM) proliferation, and vascular maturation.(*1*) In the adult heart, the epicardium retains a capacity for reactivation post-injury, recapitulating certain fetal gene programs. This has prompted recent efforts to harness its paracrine influence therapeutically to modulate inflammation, support angiogenesis, reduce fibrosis, and enhance tissue repair in the injured heart.(*1*)

Extracellular vesicles (EV) have emerged as critical mediators of intercellular communication in the heart, with their dynamic miRNA and protein cargo influencing cardiac physiology, pathology, and therapeutic repair strategies.(*3–6*) Early evidence linked pericardial fluid exosomes to epicardial-driven responses in myocardial infarction (MI) and epicardial adipose tissue-derived EVs have demonstrated impacts in various cardiac diseases.(*7–11*) Epicardial cell-secreted EVs (Epi-EVs) were also shown to exhibit effects on CM proliferation(*12*) and to undergo proteomic changes in hypoxia.(*13*) In addition, cardiomyocyte cultures were instrumental in demonstrating a cardioprotective role of endothelial-EVs during IRI.(*3*) Yet, due to the difficulty of obtaining human epicardial-myocardial samples for EV isolation and the challenges of identifying the source of EVs in multi-cellular tissue explants, characterization of the profile of epicardial cell-secreted EVs and their roles in modulating the physiology of the healthy and diseased myocardium, especially in early days after injury, remains limited.(*4*)

To address this, advanced in vitro platforms are vital for accurately capturing the nuanced effects of epicardial-EVs under physiologically relevant conditions. Existing approaches, such as self-assembled cardiac organoids derived from pluripotent stem cells or engineered heart tissues (EHTs) composed of pluripotent stem cell-derived CMs and epicardial cells, have emerged to provide valuable insight into epicardial-myocardial crosstalk.(*14–18*) Recently, our group introduced an epicardial Biowire heart-on-a-chip that relies on sequential seeding of an epicardial cell layer around pre-formed cylindrical myocardial tissue, recapitulating the developmental process of epicardial cell migration and establishing a model of simulated cardiac ischemia-reperfusion injury (IRI).(*19*) This robust human epicardial-myocardial tissue platform now makes it possible to interrogate how epicardial-EVs influence cardiac physiology and pathology.

Here, by using a physiologically relevant model of human pluripotent stem cell derived epicardial-myocardial tissue (**Fig. 1**), we investigate the composition of epicardial-derived EVs before and after EMT induction and assess their functional impact in the cardiac response to ischemic injury. These findings lay the foundation for future targeted interventions that modulate epicardial signalling and next-generation EV therapeutics aimed at enhancing heart repair.

**Figure 1.**
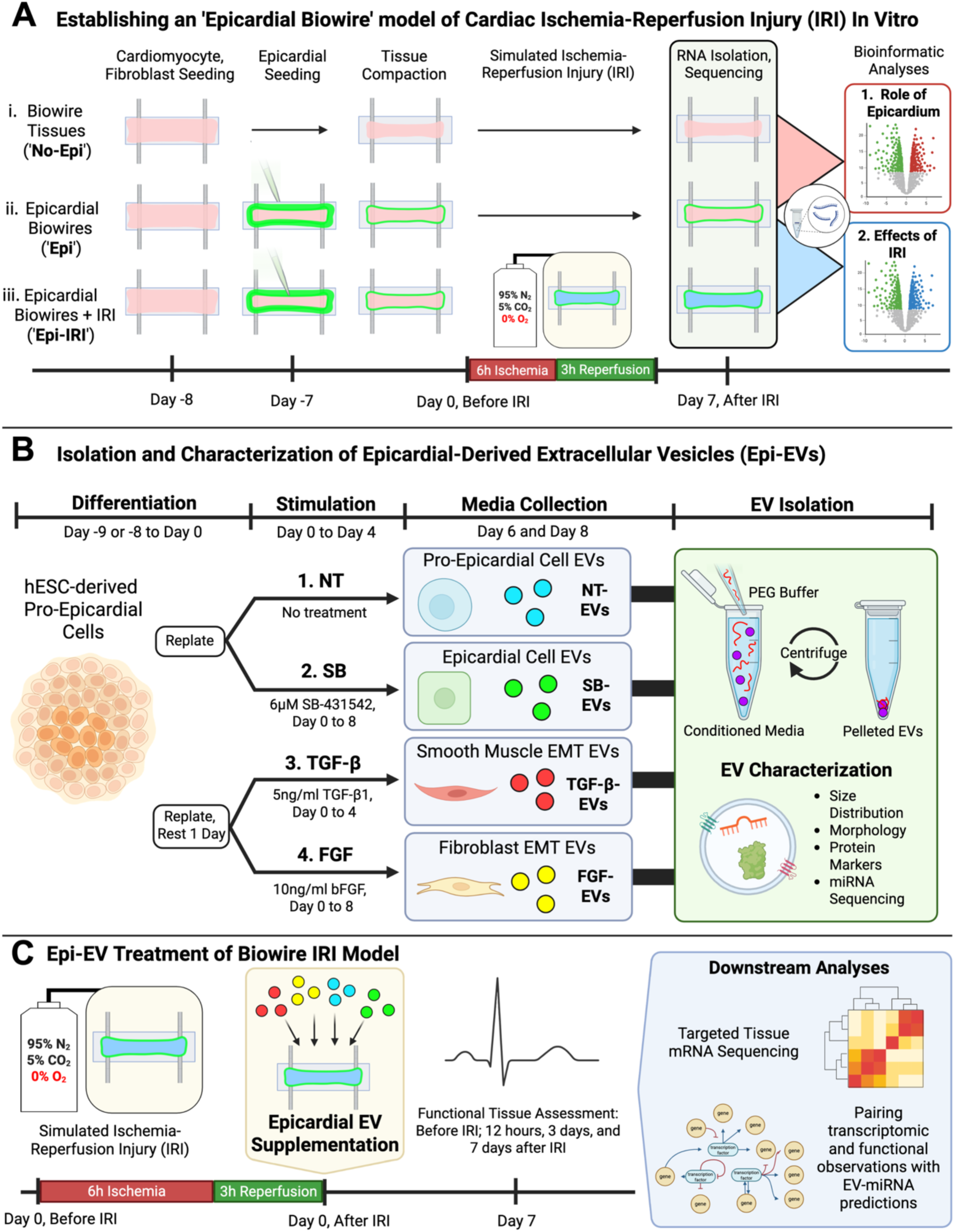
Heart-on-a-chip approach for investigating the role of epicardial cells and their EVs in cardiac ischemia-reperfusion injury (IRI). (A) Timeline for Biowire tissue seeding and injury induction. (B) Differentiation and EMT stimulation protocols were used to create four different lineages of human epicardial-derived cells from which EVs were isolated and characterized. (C) Supplementation of tissue culture media with Epi-EVs after IRI was used to assess the roles and effects of Epi-EVs in the cardiac response to injury.

## Results

Epicardial cells enhance gene expression related to extracellular matrix development and vesicle mediated transport in engineered cardiac tissues under normoxia

We adapted the ‘Biowire’ platform to fabricate engineered cardiac tissues-on-a-chip(*19–21*), incorporating a layer of epicardial cells surrounding contractile myocardial tissue to more closely mimic the arrangement of cells in the human heart and to study the effects of epicardial cells on engineered tissue phenotype via transcriptomics (**Fig.1**).

Using established protocols, we created tissues containing human induced pluripotent stem cell (hiPSC)-derived cardiomyocytes (CM) and primary human cardiac fibroblasts (CF) seeded within a collagen-Matrigel® hydrogel on the 24-well plate Biowire platform.(*21*) After one day of culture, a subset of tissues were subsequently seeded with human embryonic stem cell (hESC)-derived green fluorescent protein (GFP)-expressing epicardial cells within an identical hydrogel to create an epicardial layer surrounding the engineered myocardial tissue (**Fig. 1A(ii)**, **Fig. 2A**). In previous work published by our group with this model, functional analyses of contractility and electrical excitability did not reveal significant differences between tissues with (‘Epi’) and without epicardial cells (‘No Epi’) after two weeks of culture.(*19*)

**Figure 2.**
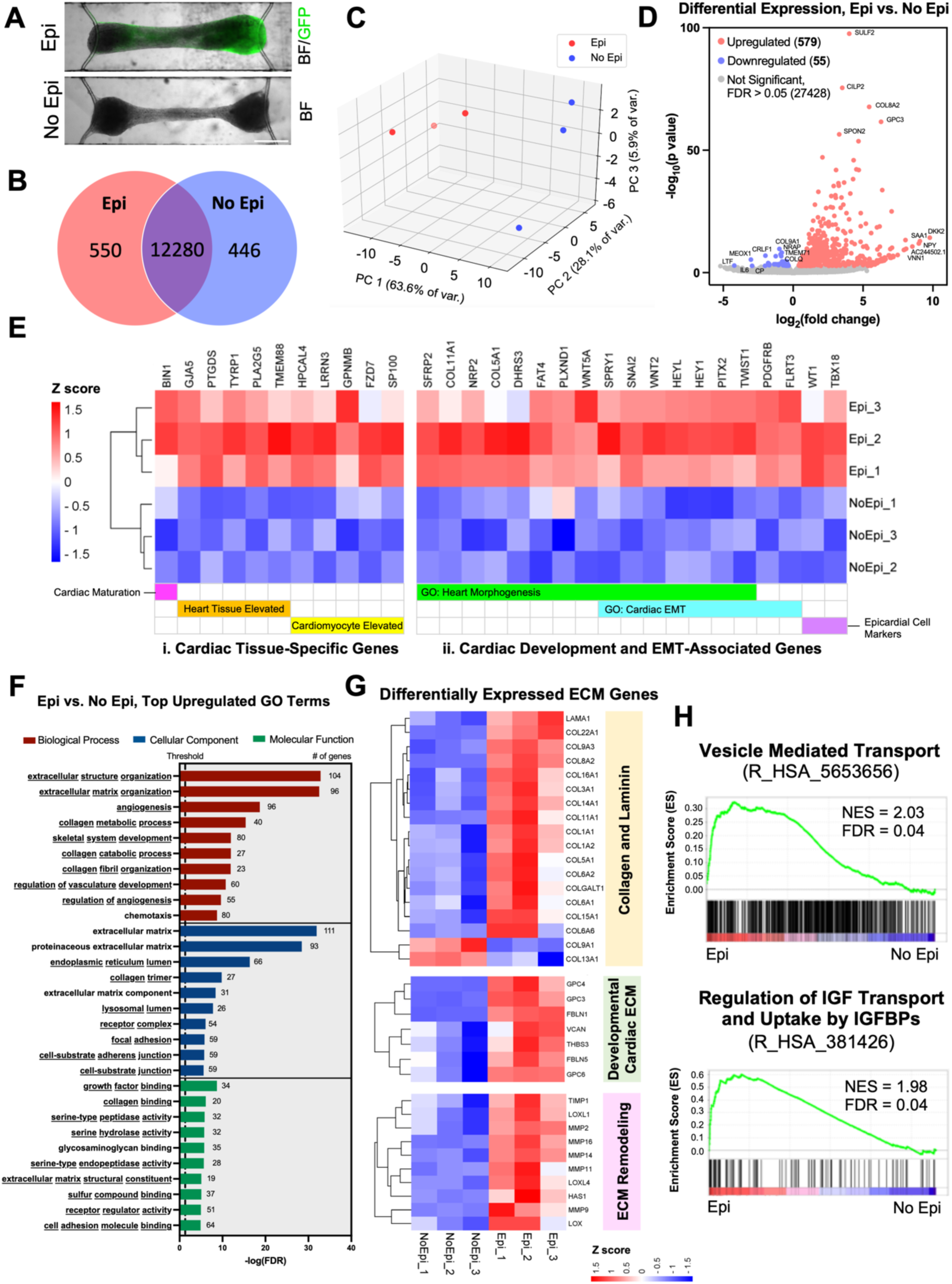
Inclusion of epicardial cells in heart-on-a-chip impacts gene expression related to cardiac development, extracellular matrix organization, and vesicle mediated transport. (A) Images of Biowire engineered cardiac tissues with GFP+ epicardial cells (‘Epi’, green, overlayed on brightfield (BF) image) and without epicardial cells (‘No Epi’) after 2 weeks of culture. Scale bar: 500µm. (B) Venn diagram depicting distinct and overlapping transcripts from mRNA sequencing of Biowire tissues with and without epicardial cells performed after 2 weeks of culture. (C) Principal component analysis (PCA) of transcriptomic data for Epi and No Epi tissue groups. (D) Volcano plot depicting significantly differentially expressed genes (DEGs) in Epi versus No Epi tissues. (E) Heatmap showing significantly upregulated genes in Epi tissues: (i) overlapping with Human Protein Atlas heart tissue/cardiomyocyte (CM)-specific genes and CM maturation genes; and (ii) associated with Epi tissue enriched gene ontology (GO) terms related to heart morphogenesis and cardiac epithelial to mesenchymal transition (EMT) alongside epicardial cell markers. (F) Top 10 significantly enriched GO terms in Epi tissues from the biological process (BP), cellular component (CC), and molecular function (MF) categories. Underlined text indicates that a given GO term is uniquely significant for Epi tissues and is not significant for No Epi tissues. (G) Heatmap showing all significant DEGs in Epi vs. No Epi tissues belonging to the categories of collagen and laminin genes, developmental cardiac ECM genes, and ECM remodeling genes. (H) Gene set enrichment analysis (GSEA) of select Reactome pathway gene sets significantly enriched in Epi tissues, including the normalized enrichment score (NES) and false discovery rate-adjusted p-values (FDR) for each gene set. For (B)-(G): n=3 for both groups; FDR<0.05 considered significant. For (H): n=3 for both groups; FDR<0.25 considered significant for GSEA.

At the two-week timepoint of culture, bulk mRNA sequencing uncovered numerous distinct and overlapping mRNA transcripts detected in tissues with and without epicardial cells (**Fig. 2B, C; Supplemental Tables 1, 2**). Differential expression analysis indicated that 579 genes were significantly upregulated and 55 were significantly downregulated with the inclusion of epicardial cells in tissues (**Fig. 2D; Supplemental Fig. S1A, Supplemental Table 3**). Filtering upregulated genes against the Human Protein Atlas list of human heart tissue- and CM-elevated genes(*22, 23*) alongside published CM maturation-related genes(*24*) revealed enhanced expression of 11 markers in tissues with epicardial cells compared to those without (**Fig. 2E(i)**), despite the expected dilution of myocardial-associated transcripts with the addition of the epicardial cell type to Biowires. These genes included bridging integrator 1 (*BIN1*) and gap junction protein alpha 5 (*GJA5*). An additional subset of 17 genes upregulated in Epi tissues was identified by their connection to significantly enriched gene ontology (GO) terms associated with heart morphogenesis and cardiac epithelial to mesenchymal transition (EMT). Two key epicardial cell markers, WT1 transcription factor (*WT1*) and T-box transcription factor 18 (*TBX18*)(*25, 26*), were also enriched in Epi tissues (**Fig. 2E(ii)**).

Globally, GO analysis(*27, 28*) of differentially expressed genes (DEGs) alluded to the downregulation of cardiac processes in Epi tissues which, by nature, included a smaller proportion of CM due to the addition of epicardial cells during tissue seeding (**Supplemental Fig. S1B, Supplemental Table 5**). However, GO analysis also suggested that numerous terms related to extracellular matrix (ECM) organization and constituents were uniquely enriched in Epi tissues (**Fig. 2F; Supplemental Table 4**).

Indeed, a search of all DEGs yielded 16 upregulated collagen/laminin genes, 7 upregulated genes known to be associated with developmental cardiac ECM, and 10 upregulated genes associated with ECM remodeling(*29, 30*). Conversely, there were only two collagen genes downregulated in Epi tissues (*COL9A1, COL13A1*) (**Fig. 2G**).

Gene set enrichment analysis(*31, 32*) (GSEA) of Reactome(*33*) pathways further indicated that vesicle mediated transport and regulation of insulin-like growth factor (IGF) transport and uptake by IGF binding proteins (IGFBPs) were among the significantly enriched pathways in Epi tissues compared to No Epi controls. Both of these gene sets were among those tied for the highest level of significance in Reactome GSEA analysis (false discovery rate-adjusted p-value (FDR)=0.04), recording strong normalized enrichment scores (NES) of 2.03 and 1.98, respectively (**Fig. 2H; Supplemental Table 6**). Collectively, these data suggest that inclusion of Epi cells promoted defined cardiac marker expression, despite overall transcript dilution, while significantly enhancing processes related to ECM development and EV transport, both critical for tissue remodelling during developmental stages and into adulthood.

### Epicardial-derived cells without and with EMT stimulation secrete EVs with commonly enriched miRNA cargo

Informed by our findings of altered gene expression and enhanced EV signaling in Epi tissues, we next aimed to isolate EVs from epicardial-derived cells to further characterize their miRNA cargo. Seeding hESC-derived pro-epicardial cells in 2D, we separately applied different culture conditions established in previous literature(*25*) to generate three lineages: mature epicardial cells (stimulated with the ACTIVIN/NODAL/TGFB inhibitor SB-431542 (SB) to complete epicardial differentiation(*25*)), those stimulated to undergo EMT with transforming growth factor-β (TGF-β) , and those stimulated to undergo EMT with basic fibroblast growth factor (FGF). Unstimulated pro-epicardial cells were also cultured as a no treatment control (NT). Immunofluorescence (IF) imaging of these cells indicated significantly increased expression of α-smooth muscle actin (α-SMA) in both EMT groups (stimulated with TGF-β or FGF), whereas TGF-β-treated cells expressed increased levels of vimentin and FGF-treated cells contained fewer zonula occludens-1 (ZO-1) positive tight junctions compared to the mature epicardial cells (**Supplemental Fig. S2**), collectively indicating progression to EMT.

Nanoparticle tracking analysis (NTA) of EVs isolated from different epicardial populations (**Fig. 3A; Supplemental Fig. S3A**) did not reveal any significant differences in particle concentration or mean size (**Fig. 3B, C; Supplemental Fig. S3B, C**).

**Figure 3.**
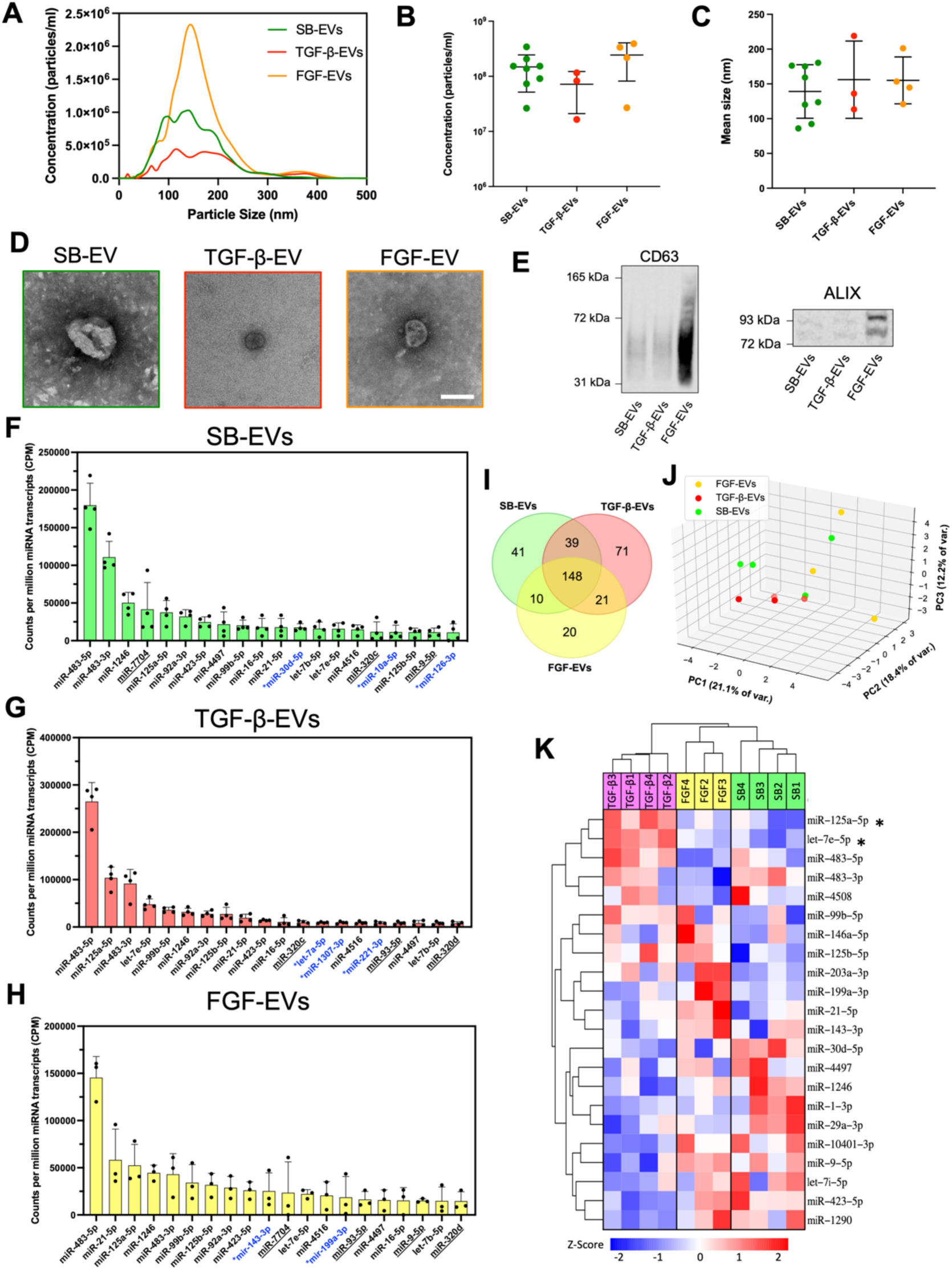
Extracellular vesicles isolated from conditioned media of epicardial cells (SB-EVs) and those undergoing EMT (TGF-β-EVs, FGF-EVs) reveal similarities in their miRNA cargo. (A-C) Nanoparticle tracking analysis (NTA) was used to quantify (A) averaged size distribution curves, (B) concentration, and (C) mean size of epicardial-EVs. (D) Transmission electron microscopy (TEM) images of epicardial-EVs. Scale bar: 100nm. (E) Western blotting detection of CD63 and ALIX in epicardial-EV isolates. (F-H) Top 20 most abundant miRNAs detected from miRNA sequencing of (F) SB-EVs, (G) TGF-β-EVs, and (H) FGF-EVs. Underlined miRNA labels indicate that a given miRNA is present in the top 20 miRNAs for two out of the three EV types, while labels with *blue text indicate that a given miRNA is unique to the top 20 miRNAs of a given group. All other miRNAs were common to the top 20 of all three EV types. (I) Venn diagram depicting distinct and overlapping miRNA cargo in epicardial-EVs. (J) PCA plot for epicardial-EV miRNA. (K) Heatmap depicting the relative expression of all miRNAs identified to have a nominal p-value<0.05 in at least one pairwise differential expression analysis comparison between the three EV types. * indicates FDR<0.05. For (A)-(C): n=8 for SB-EVs, n=3 for TGF-β-EVs, n=4 for FGF-EVs. For (B),(C): One-way ANOVA with post-hoc Tukey test; P<0.05 considered significant. For (F)-(K): n=4 for SB- and TGF-β-EVs, n=3 for FGF-EVs. Data presented as mean ± standard deviation (SD).

Transmission electron microscopy indicated the presence of particles with a characteristic cup-shaped morphology in isolates (**Fig. 3D; Supplemental Fig. S3D**), with western blotting positively detecting the presence of the EV membrane marker CD63 and the intraluminal EV marker ALIX(*34*) (**Fig. 3E; Supplemental Fig. S3E, S4**).

Isolation and sequencing of EV-miRNA cargo from each lineage of epicardial cells indicated overlap in the most abundant miRNA species present (**Fig. 3F-H; Supplemental Fig. S3F, Supplemental Tables 13-15**). Specifically, 14 common miRNAs were found within the top 20 most abundant transcripts in SB-stimulated mature epicardial cell-derived EVs (SB-EVs) and the two EMT-stimulated lineages (denoted as TGF-β-EVs and FGF-EVs). miR-483-5p was the most abundant miRNA in each EV type. miRNAs uniquely abundant in SB-EVs as compared to those secreted by EMT-stimulated cells included miR-30d-5p, miR-10a-5p, and miR-126-3p. Between the SB-EVs, TGF-β-EVs, and FGF-EVs, there were 350 total distinct and overlapping miRNA species detected (**Fig. 3I, J**). Pairwise differential expression analysis between the 3 groups of EVs nominally identified 22 miRNAs (p-value<0.05), though only two were significantly differentially expressed (FDR<0.05): miR-125a-5p and let-7e-5p, both of which were downregulated in SB-EVs compared to TGF-β-EVs (**Fig. 3K; Supplemental Fig. S5, Supplemental Table 16**).

### In Vitro Ischemia-Reperfusion Injury Perturbs Gene Expression in Epicardial-Myocardial Heart-on-a-Chip

Seeking to understand the effects of epicardial-EVs in the cardiac injury response, we first characterized IRI-induced changes to engineered tissue gene expression. One week after seeding, a subset of epicardial Biowires were exposed to hypoxic culture in an acidic ‘ischemic media’ solution followed by reperfusion in normoxia with calcium-supplemented media to induce intracellular calcium overload(*19, 35, 36*) (**Fig. 4A, B**). We previously showed that in this model, the average tissue contractility was directionally lower in tissues 12 hours after IRI and did not significantly increase over 1 week post-injury in contrast to uninjured control tissues, which exhibited increased active tension over the same period(*19*). One week after IRI, RNA sequencing from both injured (Epi + IRI) and uninjured control (Epi) tissues detected numerous distinct and overlapping mRNA transcripts between the two groups (**Fig. 4C, D**).

**Figure 4.**
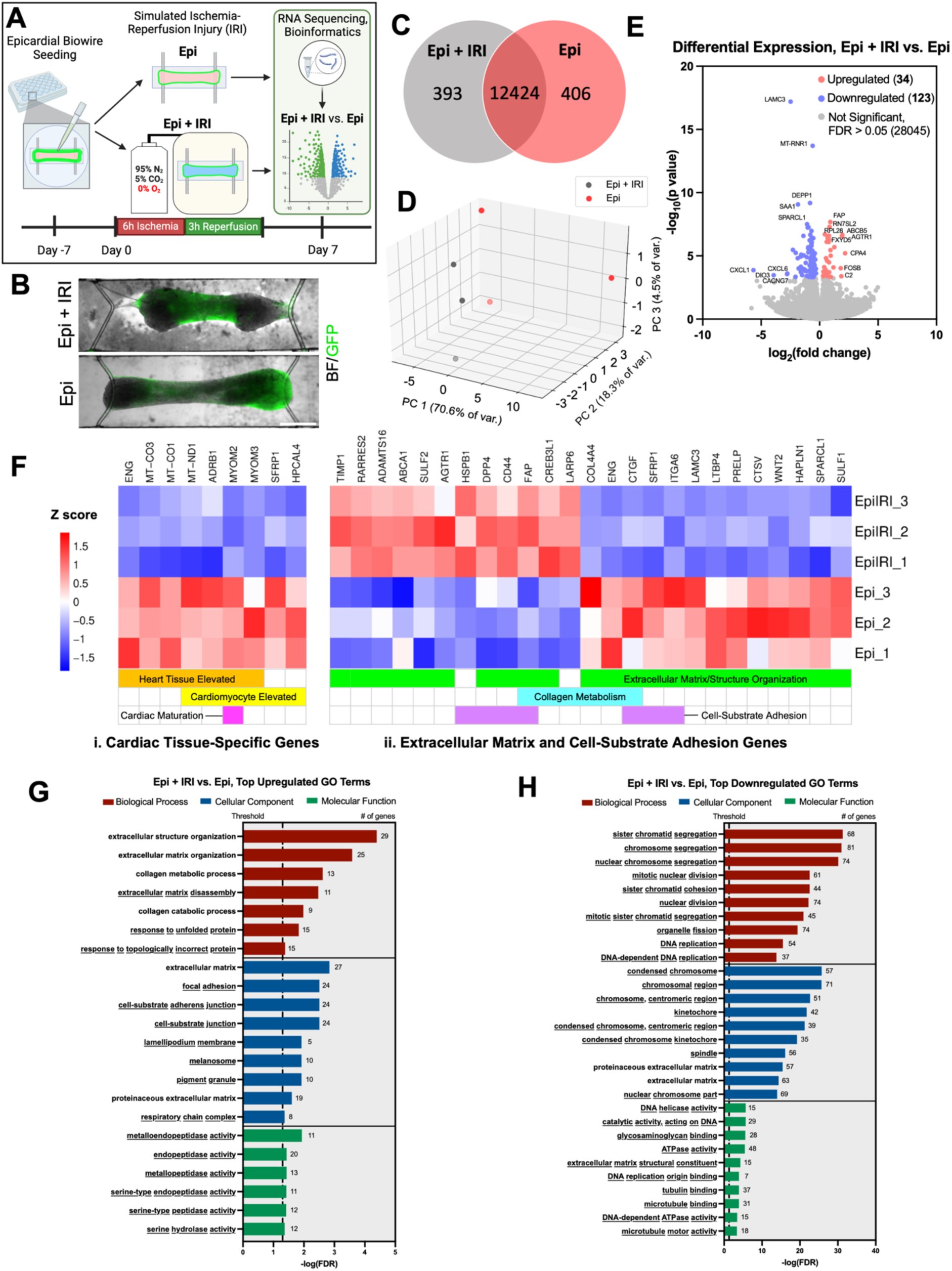
Ischemia-reperfusion injury (IRI) in epicardial heart-on-a-chip resultes in gene expression changes enhancing ECM production and remodelling while decreasing cell proliferation. (A) Schematic depicting IRI induction in engineered tissues. (B) Images of representative Epicardial Biowire tissues one week after IRI (‘Epi + IRI’) compared to uninjured controls (‘Epi’). GFP+ epicardial cells (green) overlaid on brightfield (BF) images. Scale bar: 500µm. (C) Venn diagram depicting distinct and overlapping transcripts detected in mRNA sequencing of injured versus uninjured Biowire tissues. (D) PCA plot for mRNA sequencing of Epi + IRI vs. Epi tissue groups. (E) Volcano plot depicting significant DEGs in Epi + IRI versus Epi tissues. (F) Heatmap showing significant DEGs: (i) overlapping with Human Protein Atlas heart tissue/cardiomyocyte (CM)-specific genes and CM maturation genes; and (ii) associated with enriched GO terms related to ECM organization and cell-substrate adhesion. (G, H) Top 10 significantly enriched GO terms from the BP, CC, and MF categories for DEGs (G) upregulated in Epi + IRI tissues and (H) downregulated in Epi + IRI tissues compared to uninjured Epi controls. Underlined text indicates that a given GO term is uniquely significant to the comparison where it appears and is not significantly enriched in the opposite direction. For (C)-(H): n=3 for both groups; FDR<0.05 considered significant. (A) Schematic created using Biorender.

Differential expression analysis identified 34 significantly upregulated genes and 123 significantly downregulated genes in injured tissues compared to uninjured controls (**Fig. 4E; Supplemental Table 7**). Filtering all DEGs against the Human Protein Atlas heart tissue/CM elevated genes and CM maturation genes yielded 9 matches, all of which were significantly downregulated in tissues post-IRI (**Fig. 4F(i)**). These genes included the sarcomeric M-band proteins myomesin-2 and myomesin-3 (*MYOM2, MYOM3*)(*37*). Noting the aforementioned impacts of epicardial cells on ECM gene expression, we also performed GO analysis to extract and filter DEGs associated with extracellular matrix-, collagen metabolism-, and cell-substrate adhesion-related GO terms that were enriched in injured and/or uninjured tissues (**Fig. 4F(ii)**). We observed differential expression of 25 genes from these categories, including IRI-mediated upregulation of fibroblast activation protein alpha (*FAP*), TIMP metallopeptidase inhibitor 1 (*TIMP1*)*, CD44,* angiotensin II receptor type 1 (*AGTR1*), ADAM metallopeptidase with thrombospondin type 1 motif 16 (*ADAMTS16*), La ribonucleoprotein 6 (*LARP6*), and cAMP responsive element binding protein 3 like 1 (*CREB3L1*). Downregulated genes in injured tissues included collagen type IV alpha 4 (*COL4A4*), laminin subunit gamma 3 (*LAMC3*), integrin subunit alpha 6 (ITGA6), connective tissue growth factor (*CTGF*), endoglin (*ENG*), and latent TGF-β binding protein 4 (*LTBP4*). Global GO analysis of DEGs indicated unique enrichment of extracellular matrix disassembly, cell-substrate (adherens) junction, and focal adhesion terms in injured tissues (**Fig. 4G; Supplemental Table 8**), while revealing downregulation of numerous terms related to cell division after IRI (**Fig. 4H; Supplemental Table 9**).

### Epicardial-EVs Modulate Gene Expression Following IRI in Heart-on-a-Chip

Having established the epicardial Biowire model of IRI and confirming that tissues can uptake supplemental EVs (**Supplemental Fig. S6**), we next examined whether epicardial-EVs influence the myocardial injury response. To this end, we supplemented the culture medium of epicardial Biowire tissues immediately post-injury with EVs isolated from each epicardial lineage separately (**Fig. 1C**). This revealed comparable effects across EVs isolated from cells without and with EMT on contractility and cell migration (**Supplemental Fig. S7, S8**). Since our characterization of EV-miRNA cargo also revealed significant overlap between the EVs from epicardial cells without and with EMT, we decided to narrow our experimental focus solely to EVs derived from mature epicardial cells (SB-EVs) (**Fig. 5A**).

**Figure 5.**
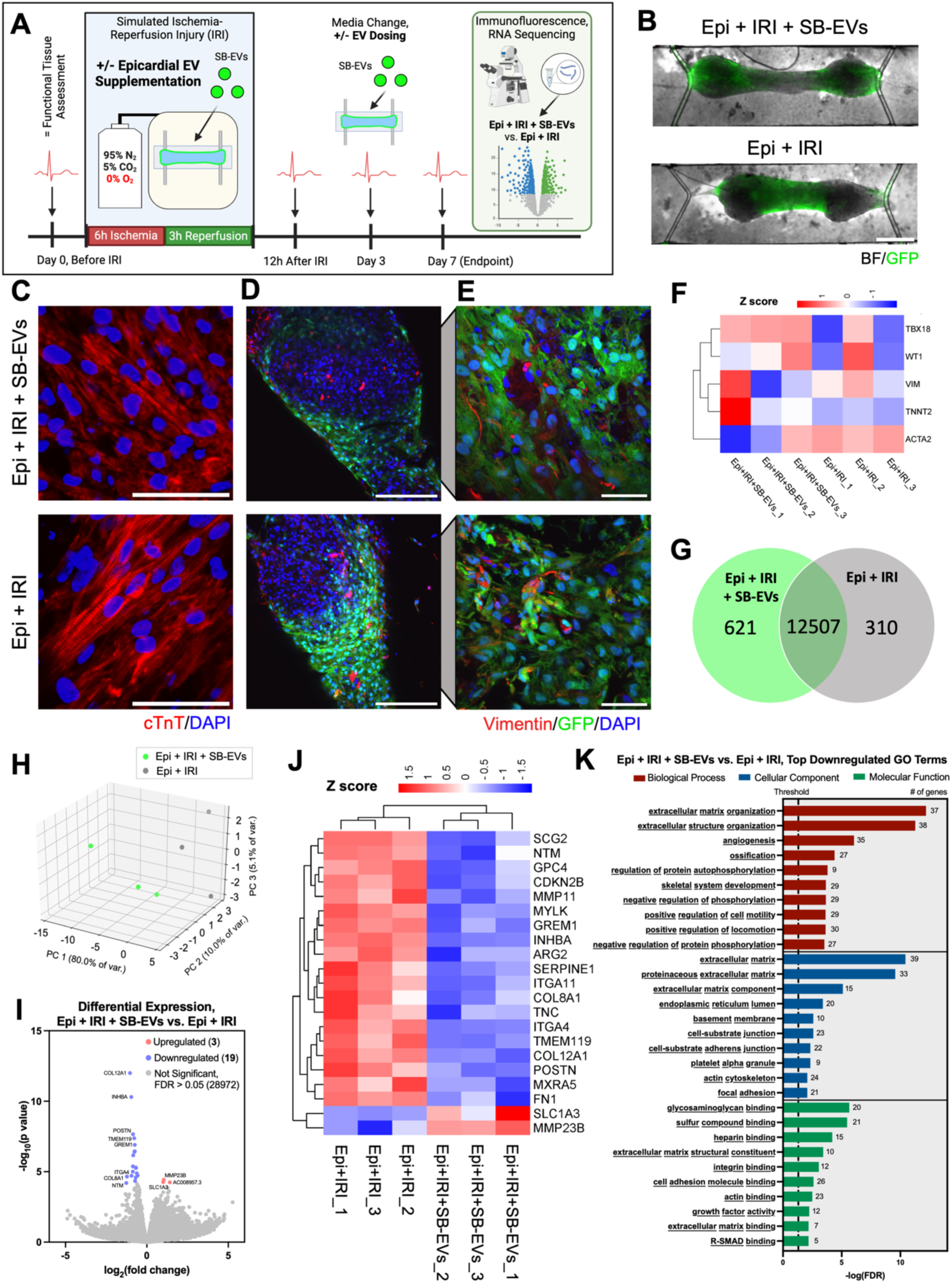
Supplementation of heart-on-a-chip with epicardial extracellular vesicles during and after IRI downregulates genes associated with ECM remodeling and cell-ECM interactions. (A) Schematic depicting SB-EV supplementation to engineered tissues and tissue characterization. (B) Images of representative Epicardial Biowire tissues supplemented with SB-EVs during and for one week after IRI (‘Epi + IRI + SB-EVs’) compared to injured controls without supplemental EVs (‘Epi + IRI’). GFP+ epicardial cells (green) overlaid on brightfield (BF) images. Scale bar: 500µm. (C) Representative IF images of cardiac troponin T (cTnT, red) in injured Biowires with and without SB-EV supplementation. Nuclei are stained with DAPI (blue). Scale bars: 50µm. (D, E) Representative IF images of vimentin (red) staining of Biowire tissues, also showing GFP+ epicardial cells (green) and DAPI-stained nuclei (blue). Tissues are depicted at (D) 20x magnification (scale bars: 200µm) and (E) 60x magnification (scale bars: 50µm). (F) Heatmap comparing relative gene expression of select key cardiac (TNNT2), epicardial (WT1, TBX18), and EMT (VIM, ACTA2) cellular markers detected in each tissue. Abbreviations: TNNT2 – cardiac troponin T; WT1 – WT1 transcription factor; TBX18 – T-box transcription factor 18; VIM – vimentin; ACTA2 - α-smooth muscle actin. (G) Venn diagram depicting distinct and overlapping transcripts, (H) PCA plot, (I) volcano plot, and (J) heatmap showing significantly DEGs for mRNA sequencing of Epi + IRI + SB-EVs versus Epi + IRI tissue groups. (K) Top 10 significantly enriched GO terms from the BP, CC, and MF categories for genes significantly downregulated in SB-EV supplemented injured tissues compared to injured controls. Underlined text indicates that a given term is uniquely significant to downregulated GO terms and is not significantly enriched in GO analysis of upregulated DEGs. For (F)-(K): n=3 for both groups; FDR<0.05 considered significant. (A) Schematic created using Biorender.

Targeted dosing of epicardial Biowires with SB-EVs did not induce significant changes in tissue contractility or excitability at one week post-injury (**Supplemental Fig. S9**). At this time point, the tissues with and without SB-EV supplementation (‘Epi + IRI + SB-EVs’ and ‘Epi + IRI’, respectively; **Fig. 5B**) both exhibited striations in cardiac troponin T (cTnT; *TNNT2*) structure (**Fig. 5C**) with the presence of GFP+ epicardial cells and vimentin (*VIM*) (**Fig. 5D, E**). Expression of cell-specific markers indicated a similar cell composition in tissues with and without SB-EV supplementation (**Fig. 5F**).

Globally, transcriptomic analysis detected numerous distinct and overlapping mRNAs in injured tissues with and without SB-EVs supplementation (**Fig. 5G, H**). Differential expression analysis revealed 19 genes significantly downregulated in injured tissues supplemented with SB-EVs, including fibronectin 1 (*FN1*), periostin (*POSTN*), collagen type XII and VII alpha chain 1 (*COL12A1, COL8A1*), integrin subunits alpha 4 and 11 (*ITGA4, ITGA11*), tenascin C (*TNC*), myosin light chain kinase (*MYLK*), matrix metallopeptidase 11 (*MMP11*), glypican 4 (*GPC4*), matrix remodeling associated 5 (*MXRA5*), neurotrimin (*NTM*), inhibin subunit beta A (*INHBA*), and serpin family E member 1 (SERPINE1). Two genes were significantly upregulated in EV-supplemented tissues: solute carrier family 1 member 3 (*SLC1A3*) and *MMP23B* (**Fig. 5I, J; Supplemental Table 10**).

In a reversal of the trend of top GO terms enriched after IRI (**Fig. 4**), GO analysis of SB-EV-supplemented tissues highlighted the unique downward enrichment of numerous terms related to extracellular matrix structure and organization, as well as cell receptor-ECM interactions (**Fig. 5K; Supplemental Table 12**). Only 11 GO terms were positively enriched in tissues supplemented with SB-EVs with marginal significance, mostly related to mRNA processing (**Supplemental Fig. S10A; Supplemental Table 11**).

The miRNA target prediction databases miRDB(*38, 39*), microT(*40*), and TargetScan(*41*) were used alongside the experimental databases miRTarBase(*42*) and TarBase(*43*) to correlate mRNA changes observed from transcriptomic analysis of SB-EV treated epicardial Biowires with specific miRNAs detected in SB-EV cargo (**Fig. 6A**). Given the role of miRNAs in post-transcriptional gene silencing, we queried the five aforementioned databases to identify candidate miRNAs predicted or experimentally validated to target and suppress any of the 19 significantly downregulated genes in SB-EV supplemented tissues. Filtering candidate miRNAs to retain only those that were within the top 20 most abundant in SB-EVs yielded a list of interactions between SB-EV miRNA cargo and their predicted mRNA targets in injured epicardial Biowire tissues (**Supplemental Table 17**). let-7b-5p had the largest total number of predicted interactions across all databases and mRNA targets with 25. This was followed by miR-30d-5p, let-7e-5p, and miR-9-5p with 24, then miR-16-5p with 20 (**Fig. 6B**). miR-483-5p, the most abundant miRNA detected in each lineage of epicardial derived EVs, only yielded one predicted miRNA-mRNA interaction (out of 194 total mapped interactions) and none that were experimentally validated. The top 5 targeted genes in our list were *COL12A1, MYLK, ARG2, FN1,* and *GPC4*, each with at least 16 combined predicted and experimentally validated miRNA interactions. (**Supplemental Fig. S10B**).

**Figure 6.**
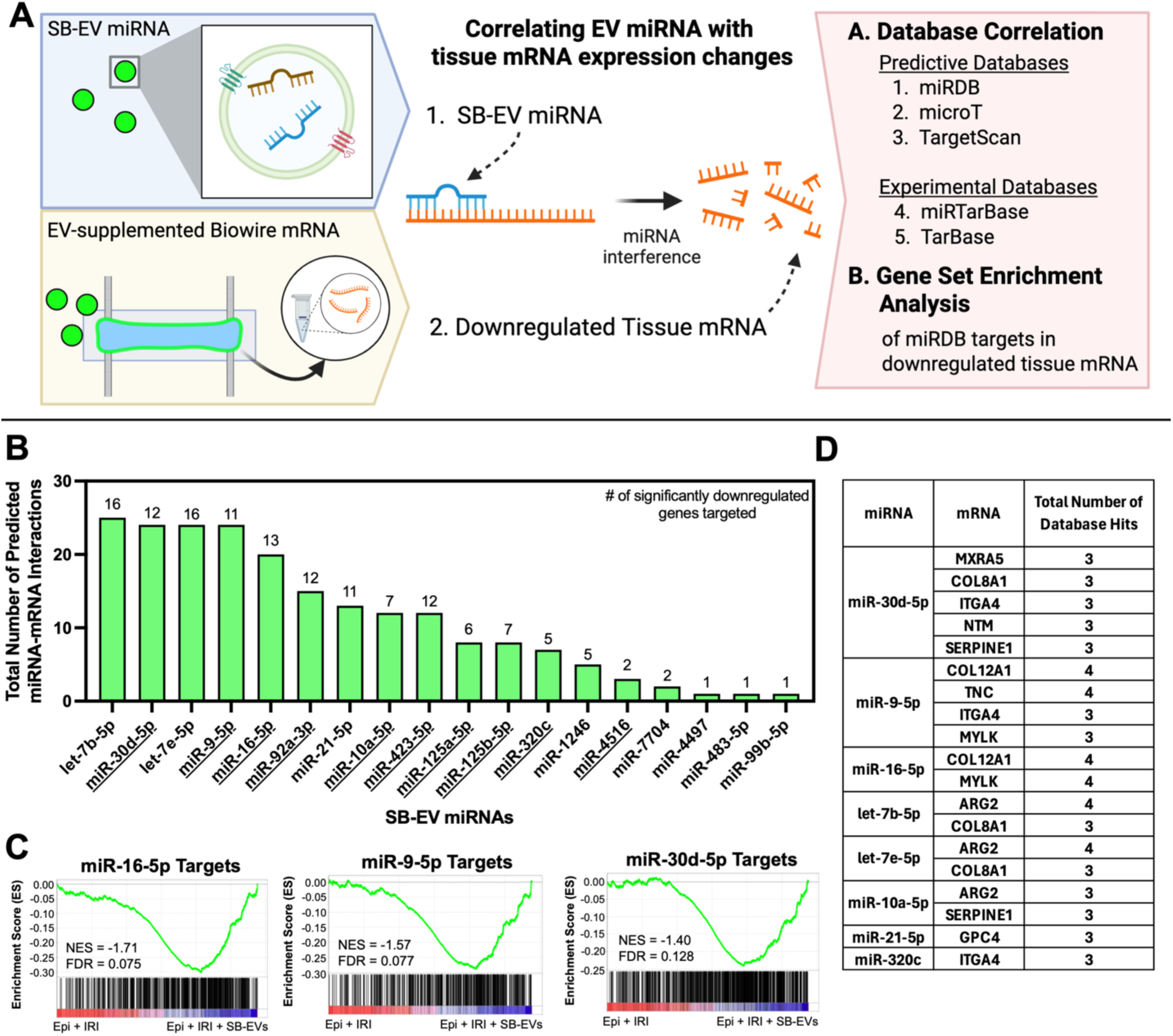
miRNAs from epicardial vesicles target key genes modulated by IRI in heart-on-a-chip. (A) Schematic depicting the theory and methodology of miRNA-mRNA correlation analyses. (B) Plot of abundant and enriched SB-EV miRNAs with the greatest total number of predicted miRNA-mRNA interactions for all Epi + IRI + SB-EV downregulated genes, combined from all five predictive/experimental miRNA databases queried. Underlined text indicates miRNAs that were also significantly enriched in miRDB GSEA (FDR<0.25). Numbers above each bar indicate the number of Epi + IRI + SB-EV downregulated genes covered by at least one predicted/experimental miRNA-mRNA interaction (out of 19 total genes). (C) GSEA of miRDB-predicted miRNA target gene sets for the top three miRNAs with the most predicted database interactions and FDR<0.25: miR-16-5p, miR-9-5p, and miR-30d-5p. Plots include the NES and FDR values for each gene set. n=3 for both groups. (D) List of correlated miRNA-mRNA interactions between abundant and enriched SB-EV miRNAs and downregulated genes in Epi + IRI + SB-EV tissues with at least 3 predictive or experimental database hits. (A) Schematic created using Biorender.

We also performed GSEA on miRDB-predicted targets in the transcriptomic data of EV-supplemented tissues as a secondary method to filter and refine the target prediction analysis, identifying miR-16-5p, miR-9-5p, and miR-30d-5p as the top three miRNA species with the largest number of total predicted miRNA-mRNA interactions in addition to significant gene set enrichment (**Fig. 6C; Supplemental Table 18**).

Finally, we sorted specific miRNA-mRNA interaction pairs by the total number of databases they appeared in (‘database hits’) to produce a short list of 25 highest confidence miRNA-mRNA interactions with 3 or more database hits that were relevant to the SB-EV-supplemented cardiac tissues (**Fig. 6D**; **Supplemental Fig. S10C**). This short list included 8 different SB-EV miRNAs targeting 10 of the 19 significantly downregulated genes in SB-EV supplemented tissues. Two different members of the let-7 family of miRNAs appeared in the short list due to interactions with arginase 2 (*ARG2*) and *COL8A1*. miR-30d-5p had five unique appearances on the short list (for interactions with *MXRA5, COL8A1, ITGA4, NTM,* and *SERPINE1*) while miR-9-5p had four (*COL12A1, TNC, ITGA4,* and *MYLK*) and miR-16-5p had two (*COL12A1* and *MYLK)*.

Collectively, these data prove that SB-EVs carry miRNA that could alter and suppress matrix production after IRI in the epicardial-myocardial heart-on-a-chip model.

## Discussion

The critical role of epicardial paracrine signaling in cardiac development and its potential in therapeutic repair have been well established.(*2*) However, the specific composition and functional contributions of EVs derived from epicardial cells, in the absence or presence of EMT, to such processes remain largely underexplored. Human in vitro models that recapitulate epicardial-myocardial interactions offer distinct advantages for such studies due to the defined cellular composition and facile external supplementation of EVs.

Here, we provide a Biowire heart-on-a-chip model, consisting of pluripotent stem cell derived cardiomyocytes and fibroblasts in the core and a layer of epicardial cells on the surface, as a useful tool to study gene expression differences resulting from IRI and Epi-EV supplementation. The current study provides the following important contributions: 1) we defined the change in transcriptome of human pluripotent stem cell derived hearts-on-a-chip with the inclusion of epicardial cells, pointing to the importance of EV signaling; 2) we detected compelling similarity in the miRNA content of EVs from mature epicardial cells and EMT-stimulated lineages; 3) we defined the impact of epicardial cell inclusion on transcriptomic changes shortly after IRI; and 4) we showed that Epi-EV supplementation did not result in inotropic effects or changes in epicardial cell migration within the tissue over the short term (7 days), yet it powerfully modulated the transcriptome of injured epicardial-myocardial hearts-on-a-chip, possibly affecting long term remodelling and paving the way to future therapeutic development.

Inclusion of epicardial cells in cardiac tissues enhanced important markers of maturation (gap junction and T-tubule transcripts(*44, 45*)) under normoxic conditions, consistent with previous studies.(*14*) In agreement our previous observation of epicardial cell migration from the outer surface to the interior of tissues post-seeding(*19*), transcriptomic analysis further highlighted the enhancement of a developmental cardiac phenotype in epicardially coated tissues, with GO terms for heart morphogenesis, cardiac EMT processes, and key associated genes solely enriched in tissues containing epicardial cells. These included the EMT markers *TWIST1*(*46*) *and SNAI2*(*47*) that were also identified in self-organizing stem cell-derived epicardioids.(*17*)

Overall, we observed compelling conservation of miRNA cargo between mature epicardial-EVs and those derived from EMT-stimulated lineages. Availability of human pluripotent stem cell derived epicardial cells enabled these studies, addressing difficulty with sampling primary human epicardial cells, especially in a quiescent state not affected by EMT. Interestingly, the hESC-derived epicardial as well as EMT-stimulated Epi-EV types were relatively abundant in miR-483-5p and -3p transcripts, not previously reported in primary human epicardial-EVs and present at low levels in mouse epicardial cell line EVs.(*12*) Abundant expression of miR-21-5p in all of the EVs from epicardial lineages was conserved in published data from primary human inactive and activated Epi-EVs.(*12*) This miRNA has previously been shown to exert positive effects on CM contractility and calcium handling in 3D engineered cardiac tissues.(*48*) Various members of the let-7 family of miRNAs, as identified in the epicardial derived EVs, were also reported in primary human Epi-EVs.(*12*) The importance of let-7 miRNAs in regulating cardiac vasculogenesis and homeostasis has been implicated by numerous studies(*6, 49, 50*), as have direct effects of let-7s on CM metabolism, cardioprotection, and disease.(*50, 51*) Such examples illustrate potential benefits of epicardial-myocardial EV signalling on enhancing physiological cardiac phenotype in vitro. miR-30d-5p, unique to the top 20 most abundant miRNAs in mature epicardial-EVs (SB-EVs) compared to the EMT-stimulated lineages, has been identified as modulator of the cardiac ischemic response in vitro and in vivo, reducing infarct size, CM apoptosis, and myocardial fibrosis while enhancing tissue contractility when overexpressed.(*52, 53*)

Epicardial-myocardial heart-on-a-chip uniquely enabled us to define the transcriptome of human tissues shortly (7 days) after IRI. These early changes are poorly understood yet critically important for treatment of disease, but they are difficult to interrogate in a human in vivo. Post-injury downregulation of heart tissue- and cardiomyocyte-elevated genes, including two sarcomeric myomesin proteins, reflected phenotypic impacts of IRI aligned with our previous observation of impaired tissue contractile maturation.(*19*) Differential reorganization of ECM was also apparent after IRI. Upregulation of *FAP, TIMP1, CD44, AGTR1,* and *ADAMTS16* reflected conditions of enhanced fibroblast activation and early ECM remodeling in injured tissues.(*54–59*) Reductions in *COL4A4, LAMC3,* and *ITGA6* were consistent with possible disruption of the cardiac basement membrane at this stage of remodelling(*60, 61*), with increased expression of collagen I regulators *LARP6* and *CREB3L1* supporting interstitial collagen deposition to come.(*62–64*)

Tissues supplemented with SB-EVs displayed a broadly apparent reversal in the trend of GO terms enriched after injury, exhibiting significant downregulation of numerous ECM-related terms. This trend was supported by an examination of specific DEGs.

Overall, our results indicated notable changes in tissue gene expression induced by externally supplemented epicardial-EVs in an in vitro model of cardiac IRI, including genes associated with reduced injury-associated ECM deposition, modulated fibroblast activation, and suppressed cell-ECM interactions. These phenotypic effects are supported by predictive and mechanistic data in literature linked to the cardioprotective role of the epicardial-EV enriched miR-30d-5p(*52*), with potential contributions of other species (including miR-9-5p, miR-16-5p, and the let-7 family) motivating future studies focused on mechanistic validation.

### Limitations and Future Work

The need for controlled and reproducible production of Epi-EVs in vitro motivated us to isolate and characterize EVs from stable 2D epicardial cell cultures obtained from a well-defined differentiation protocol. While this provided a simple way to characterize Epi-EVs, the complexity of epicardial-myocardial cross-talk in the native heart alongside the inherent diversity of epicardial cells derived from multi-faceted EMT activation pathways during injury means that the phenotype of epicardial cells and thus, the profile of miRNAs sorted into secreted EVs, may be impacted and requires further investigation. We note that the IRI model, used here and optimized in previously published work(*35, 36*) to induce physiological and reproducible CM death in 3D engineered cardiac tissues, was applied as a baseline perturbation of tissue physiology from which to derive new insights into Epi-EV functionality. In effectively taking a single ‘snapshot’ at day 7 post-IRI, it is difficult to assess in detail the temporal transitions between acute injury and remodeling phases. Longer-term studies (6-8 weeks) may be necessary to address the ultimate functional effect of Epi-EVs on heart-on-a-chip function and matrix deposition.

We emphasize here the contextual importance of miRNA-mediated regulation, where changing concentrations or synergistic interactions with other miRNAs can lead to diverging outcomes within complex signalling networks like the injured heart.(*65, 66*) Follow-up investigations with transfection of specific epicardial-EV miRNAs in physiologically relevant cardiac tissues are required to produce detailed mechanistic insights into their specific functionality in cardiac physiology and injury response, while addressing biases inherent to our predictive analysis techniques. While we focused our analyses here on epicardial-EV miRNA cargo, EV-associated proteins represent another way by which EVs can regulate tissue physiology and will be another important consideration in fully defining the role and mechanisms of epicardial-EV signalling in the heart moving forward.(*4, 13*)

## Conclusion

In this study, we demonstrated that epicardial cells significantly impact gene expression when incorporated in engineered cardiac tissues in vitro, with enriched vesicle mediated transport reflecting a key outcome of epicardial-myocardial interactions. We also performed a novel characterization of the miRNA profile of human pluripotent stem cell-derived epicardial cells in comparison to two of their EMT-stimulated lineages. While possessing limited influence on cardiac contractility, epicardial-EVs notably modulated genes related to reduction of injury-associated ECM deposition, fibroblast activity, and suppressed cell-ECM interactions after simulated cardiac IRI. These phenotypic effects were supported by direct correlation with key EV-miRNA cargo and, in the case of miR-30d-5p, aligned with previously reported mechanisms. By characterizing the profile of EVs secreted by epicardial cells and investigating their effects in an in vitro model of cardiac injury, we generated new insight showing that epicardial-EVs can tangibly modulate cardiac tissue physiology post-IRI. We find a compelling case for further consideration of Epi-EVs in developing a better understanding of epicardial-myocardial cross talk in the healthy and diseased heart alongside the opportunity to harness or amplify potential cardioprotective and pro-reparative benefits of Epi-EVs and their miRNA cargo in future therapeutics.

## Acknowledgements

Schematics were created using Biorender as indicated in figure captions. We acknowledge The Hospital for Sick Children’s Structural & Biophysical Core Facility for their equipment and support with NTA, as well as the Microscopy Imaging Lab at the University of Toronto and the Advanced Optical Microscopy Facility at the University Health Network for imaging equipment access. Special thanks to Chuan Liu, Marie-Jo Abdul-Hay, Dana Park, and James Ryan Smith for data analysis support; to Trevor Nash, Bohao Liu, and Prof. Gordana Vunjak-Novakovic for advisory support; and to Prof. Gordon Keller for generously providing pro-epicardial cells. K.T.W. was supported by an Ontario Graduate Scholarship. This work was funded by the Natural Sciences and Engineering Research Council of Canada (NSERC) Discovery Grant (RGPIN 326982-10) and Canadian Institutes of Health Research (CIHR) Foundation Grant (FDN-167274), Canada Foundation for Innovation/Ontario Research Fund grant 36442. M.R. was supported by Canada Research Chairs.

## Author Contributions

K.T.W., D.B., and M.R. designed the study. Q.W. designed and fabricated Biowire culture plates. K.T.W., D.B., and S.P. performed Biowire tissue seeding. K.T.W. and D.B. performed RNA extraction, EV isolation, miRNA extraction, IRI induction, and functional tissue analyses. K.T.W. performed NTA, WB, RNA/miRNA bioinformatics analyses, and epicardial cell staining. I.F. differentiated epicardial cells and advised on cell culture and experimental design along with G.K. L.F.J.V. analyzed functional data. S.Y. assisted with epicardial cell immunofluorescence analysis. R.K. and S.O. performed Biowire tissue immunostaining and imaging. D.A. assisted with confocal imaging of cells. K.T.W. prepared figures and wrote the manuscript, which was edited by M.R, D.B., and S.O. M.R. obtained funding for the study. All authors read and approved the manuscript.

## Conflict of Interest

M.R. is an inventor on an issued US patent describing the Biowire platform, which is licensed to Valo Health, and receives royalty income from this patent. M.R. and Q.W. are inventors on patent applications covering nanocomposite micowires used here.

## Data Availability Statement

The authors declare that the main data supporting the findings of this study are available within the manuscript, its supplementary data files, and the publicly accessible GEO repository (for EV-miRNA and tissue-mRNA sequencing data; accession numbers GSE300517 and GSE300518, respectively). Additional data are available from the corresponding author upon reasonable request.

## Materials and Methods

### Biowire plate fabrication

24-well Biowire plates were fabricated as described in detail previously.(*19, 21, 67*) Briefly, soft lithography was used to create a mould for base plates with repeating rectangular tissue microwells and grooves to house polymer microwires. Hot embossing was used to transfer the base plate mould pattern to polystyrene sheets and to bond carbon electrodes into the base plate to enable electrical stimulation of tissues.

Thermoplastic elastomer(TPE)/quantum dot (QD) nanocomposite ink was prepared and used to print autofluorescent 3D microwires onto polystyrene base plates which were then bonded to 24-well bottomless plate upperstructures using polyurethane glue. A microscale mechanical tester was used to generate force-displacement calibration curves for microwires as described.(*19, 21, 67*) Assembled Biowire plates were filled with 0.2µm filtered 70% ethanol for 2 hours, rinsed with sterile phosphate buffered saline (PBS), air dried in the biosafety cabinet (BSC) overnight, treated with 5% w/v pluronic acid in distilled water for several minutes, then air dried again before tissue seeding.

### Cell culture and differentiation

Epicardial cells and GFP+ epicardial cells were differentiated from hES2 and hES2-GFP cell lines following a protocol outlined extensively in previously published reports.(*19, 25*) Briefly, embryoid body formation (day 1-4) followed by monolayer culture (day 4-8/9) was used to produce pro-epicardial cells as described, which were dissociated and cryopreserved on day 8 or 9 of differentiation.(*19, 25*)

To prepare epicardial cells Biowire seeding, cryopreserved GFP+ pro-epicardial cells were thawed and plated at a density of 200,000 cells per well on 6 well plates pre-coated with 0.1% w/v porcine gelatin (MilliporeSigma, G1890) in PBS. Cells were then cultured for 16-20 days (until confluent) in Stempro34 culture medium supplemented with 1% pen/strep, 1% l-glutamine, 50 µg/ml ascorbic acid, 0.026% (v/v) monothioglycerol (MTG), and 6 µM SB-431542 (SB; dissolved in DMSO (18 mM) MilliporeSigma, S4317-5MG). Prior to epicardial tissue seeding, cells were dissociated with 1 mg/ml collagenase B for 1 hour then 0.4% (v/v) TrypLE at 37 °C for 3 min, gently pipetted, diluted with media, pelleted at 300g for 5 min, washed with media, pelleted again, then maintained on ice.

To prepare epicardial cells for EV isolation, cryopreserved non-GFP pro-epicardial cells were seeded on 6 well plates using the same density and protocol as described above. Four different culture stimulation conditions were applied to different subsets of pro-epicardial cells. The ‘SB’ epicardial cell group was cultured in the same supplemented StemPro media formulation as outlined above from the time of plating (day 0) for the full duration of culture. Cells in the ‘TGF-β’ stimulation group were grown in the same supplemented StemPro media, but with no SB-431542 and with the addition of 5 ng/ml TGF-β1 (PeproTech, 100-21-10UG) starting one day after plating (considered day 0) for 4 days total. The ‘FGF’ stimulation group was grown in supplemented StemPro media with no SB-431542 and with the addition of 10 ng/ml bFGF (R&D Systems, 233-FB-025) starting one day after plating (day 0) for the full duration of culture. ‘No treatment’ (NT) cells were cultured in supplemented StemPro media with no SB-431542 and no other additives from the time of plating (day 0) for the full duration of culture. Culture media was changed every 2 days. On day 4, culture media (following the formulations specified above for each stimulation group) was ultrafiltered via centrifugation in Amicon Ultra-15 30kDa centrifugal filters (MilliporeSigma, UFC903008) at 4000g for 30 minutes at room temperature to deplete supplemented media of background EVs. Cells from each stimulation group were cultured in their respective formulations of ultrafiltered media for two 48-hour periods (day 4 to day 6 and day 6 to day 8). Conditioned media collected from cells on days 6 and 8 was frozen at -20°C until EV isolation. On day 8, cells from each stimulation group were fixed in 4% paraformaldehyde for 15 minutes at room temperature prior to storage in PBS for subsequent immunostaining for epicardial and EMT markers.

Human ventricular cardiac fibroblasts (CFs) (Lonza, CC-2904) were cultured in Fibroblast Growth Medium 3 (PromoCell, C-23025) with 1% 10,000 U/ml penicillin/streptomycin. Prior to Biowire seeding, CFs were treated with 0.05% trypsin for 4 min at 37 °C, neutralized with media, pelleted via centrifugation at 300 g for 5 min, resuspended in media, counted, then temporarily kept on ice until seeded.(*19*)

Frozen iCell human cardiomyocytes (Fujifilm Cellular Dynamics, 01434) were thawed according to the manufacturer’s instructions and diluted with Induction 3 Medium (I3M) consisting of supplemented Stempro34 media with 1% pen/strep, 213 μg/ml 2-Phospho-L-ascorbic acid trisodium salt (MilliporeSigma, 49752), 150 μg/mL transferrin (MilliporeSigma, T8158), 20 mM HEPES, and 1%GlutaMAX. Diluted cells were pelleted at 300g for 5 min, resuspended in media, counted, then temporarily kept on ice until seeded.(*19*)

### Biowire tissue seeding

Biowire tissues were seeded following a previously published protocol.(*19, 20*) Briefly, a hydrogel solution consisting of 10% v/v Medium 199 10x, 10% v/v 2.2 mg/ml NaHCO3, 0.6% v/v 1M NaOH, distilled water (volume adjusted depending on collagen stock concentration), 3 mg/ml collagen I (Corning, 354249), and 15% v/v GFR Matrigel was prepared. Hydrogel was adjusted to pH 7 using minute amounts of 1M NaOH and kept on ice.

CMs and CFs prepared and counted as outlined above were mixed at a ratio of 90,000 CMs and 10,000 CFs per tissue seeded then centrifuged at 300g for 5 min with careful removal of supernatant from the cell pellet. Pelleted cells were resuspended in hydrogel at a density of 50,000 cells/µl using gentle but thorough pipetting then placed on ice. 2 μl of cell-hydrogel suspension was pipetted into each tissue microwell of a Biowire plate then incubated for 10 min at 37 °C to facilitate gelation. 1.5 ml of I3M was added to tissue wells before incubation at 37 °C.

For Biowires containing epicardial cells, tissues seeded with CM and CF were prepared as outlined above and cultured for 24 hours before epicardial cell seeding. Hydrogel was prepared as before and counted GFP+ epicardial cells were pelleted at 300g for 5 min for careful removal of supernatant before suspension in hydrogel at 50,000 cells/µl on ice. Media from one-day-old CM/CF Biowires was carefully removed from tissue wells and the small space surrounding partially compacted tissues and 1µl of epicardial cell-hydrogel suspension was pipetted around tissues in microwells to surround partially compacted cardiac tissues. Tissues were incubated at 37°C for 10 min for gelation before the addition of 1.5 ml I3M. Media was replaced with 1 ml fresh I3M on days 5 and 7 after seeding.(*19*)

### Tissue mRNA isolation and sequencing

Total RNA was isolated from Biowire tissues 15 days after the initial CM/CF seeding (which came 14 days after epicardial cell seeding and 7 days after IRI, when applicable, such that all groups were harvested after the same culture duration) using the PicoPure RNA Isolation Kit (ThermoFisher, KIT0204).

Media was removed from Biowire tissue wells, and tissues were washed with PBS. A P10 pipette tip was used to gently scrape and detach the sides of each tissue from TPE wires without disturbing bulk tissue structure. Detached tissues and excess PBS were withdrawn from wells using a P1000 pipette and placed separately in microcentrifuge tubes. PBS was gently removed from the microcentrifuge tube using a pipette, allowing the tissue to remain stuck to the tube wall. 100µl of extraction buffer from the RNA isolation kit was added to the microcentrifuge tube and tissues were lysed by vigorous pipetting and vortexing.

Column-based total RNA extraction, washing, and elution was performed according to the kit manufacturer’s protocol, using 13µl of elution buffer to elute isolated RNA. Isolated RNA was stored at -80°C before shipping on dry ice to Novogene Corporation Inc. for RNA purification, quality control (QC), library preparation, sequencing, and bioinformatics services. Briefly, total RNA was purified using the RNeasy MinElute Cleanup Kit (Qiagen, 74204) according to the manufacturer’s protocol then quantified on an Agilent 5400 Bioanalyzer. For library preparation, mRNA was purified from total RNA using polyT oligo-attached magnetic beads. mRNA was fragmented, random hexamer primers were used for first strand cDNA synthesis, then dTTP for the second strand (non-directional library).(*68*) This was followed by end repair, A-tailing, adapter ligation, size selection, amplification, and purification. Libraries were quantified using a Qubit and real-time PCR as well as a Bioanalyzer for size distribution, pooled, then sequenced on a NovaSeq X Plus Series sequencer (Illumina) at a read length of 2x150 b.p. and an average depth of 20M reads. Fastq files from sequencing were cleaned up via in-house scripts for adapter trimming, poly-N trimming, and removal of low quality reads to produce data for downstream bioinformatic analyses. Tissue mRNA sequencing data was deposited in the GEO repository (accession: GSE300518).

### Downstream bioinformatic analysis of tissue mRNA sequencing data

Alignment, counting, differential expression analysis, GO, Reactome, and GSEA were performed in-house at Novogene Corporation Inc. Briefly, Hisat2 v2.0.5(*69*) was used to build the index of, and align cleaned reads to, the reference genome (GRCh38).

Mapped reads were counted using featureCounts(*70*) v1.5.0-p3 then converted to FPKM. The DESeq2 R package (1.20.0)(*71*) was used for differential expression analysis(*72*). FDR<0.05 was considered significant. The clusterProfiler(*73*) R package was used for downstream biological enrichment analyses with gene length bias corrected, including GO enrichment analysis(*28*) and Reactome(*33*) pathway analysis (FDR<0.05 considered significant for each). For each pairwise GO enrichment comparison between groups, the top 10 most significant upregulated GO terms (BP, CC, or MF) were underlined if they were not significant (FDR>0.05) in the list of downregulated GO terms (and vice versa). The Broad Institute web-based tool was used for GSEA(*32*) (https://www.gsea-msigdb.org/gsea/index.jsp; Accessed March 2024) of the Reactome dataset (FDR<0.25 considered significant).

Principal component analysis of raw counts (from featureCounts) was performed using PCAGO(*74*) (normalization using DESeq2 rlog transformation) and plotted using SRPlot(*75*). Heatmaps were also generated on SRPlot using rlog normalized counts from DESeq2 analysis performed on the Galaxy(*76*) platform, with complete Euclidean clustering of rows, columns, or both as indicated in figures.

The human protein atlas(*22, 23*) (https://www.proteinatlas.org/; Accessed December 2024; version 24.0) was used to download lists of cardiac tissue- and cardiomyocyte-specific genes from the following categories: heart enriched, heart elevated, heart group enriched, heart muscle cell type enriched (very high, high, and moderate), cardiomyocyte cell type enriched, cardiomyocyte group enriched, and cardiomyocyte cell type enhanced. These lists were each cross-referenced with significantly differentially expressed genes in sequenced tissues to highlight differentially expressed cardiac genes in Biowire tissues for heatmapping. For the custom heatmap of cardiac development and EMT genes, plotted genes were selected from the overlap of Epi upregulated genes and the gene lists associated with all Epi vs. No Epicardial Biowire significantly enriched GO terms containing the keywords ‘cardiac/heart’ AND morphogenesis, development, EMT, or formation. For the custom heatmap of Epi vs. No Epi ECM genes, all significantly differentially expressed genes (up or downregulated) from the following families were plotted: all collagen and laminin genes, developmental cardiac ECM genes (based on refs(*29, 30*)) and ECM remodelling genes (based on refs(*29, 30*)). For the custom heatmap of Epi + IRI vs. Epi tissues, plotted genes were selected from the overlap of significant DEGs in tissues (up or down) and the gene lists associated with all significant (up and/or downregulated) GO terms containing the following keywords: extracellular matrix/structure; collagen metabolic/catabolic process; ‘focal/cell-substrate’ AND ‘adhesion/junction’.

### Epicardial cell immunofluorescence imaging

Fixed epicardial cells from the four different stimulation groups (NT, SB, TGF-β, FGF) prepared as described above were imaged in brightfield on an optical microscope, briefly washed three times with PBS, then permeabilized (except for ZO-1 stained cells, which were not permeabilized before staining) with 0.25% Triton-X for 15 minutes at room temperature with gentle shaking. Cells were briefly washed three times with TBS then blocked for one hour at room temperature with 10% normal goat serum (NGS; ThermoFisher Scientific, 50062Z) with gentle shaking. Primary antibody staining was performed by incubating cells in 10% NGS overnight at 4°C with the addition of one of the following antibodies: anti-ZO-1 (ThermoFisher Scientific, 61-7300, 5µg/ml, rabbit polyclonal), anti-WT1 (Abcam, ab8990, 1:100, rabbit monoclonal), or co-stained with anti-vimentin (Abcam, ab8978, 5µg/ml, mouse monoclonal) and anti-α-SMA (Abcam, ab5694, 1:200, rabbit polyclonal). Cells were briefly washed three times with TBS before secondary antibody staining in 10% NGS for 1 hour at room temperature with gentle shaking and the addition of either goat anti-rabbit AF647-conjugated secondary (for ZO-1, WT1, α-SMA; Invitrogen, A21245, 1:500) or goat anti-mouse AF488-conjugated secondary (for vimentin; Abcam, ab150113, 1:500). Cells were briefly washed three times with TBS before incubation with DAPI (Invitrogen, D1306, 1:2000 in TBS) for 5 minutes at room temperature with gentle shaking. DAPI solution was then replaced with TBS before imaging.

Cells were imaged on an Olympus IX81 fluorescent microscope. Fiji software was used for automatic counting of DAPI+ nuclei number and coverage area. The integrated density of raw, unadjusted images was measured to quantify WT1 signal before normalizing to nuclei area. The fraction of cells with ZO-1 positive borders was manually counted. Raw integrated density of vimentin signal in unadjusted images was normalized to the number of cells in each image. The fraction of cells positive for α-SMA was manually counted.

Five regions were randomly selected and imaged for each sample (technical replicates), with each of their quantifications averaged together to produce a single biological replicate value. Three independent batches of stimulated cells were produced, with each batch containing one biological replicate from each stimulation group. Before plotting, all quantified data points described above for each marker were normalized to the corresponding ‘NT’ sample from the same batch of cells to highlight the relative differences in markers between groups.

### Epicardial-derived EV isolation

Ultrafiltered conditioned media collected separately from each group of stimulated epicardial cell cultures on days 6 and 8 and stored at -20°C as described above was thawed and used for EV isolation according to a previously published protocol.(*6*) Briefly, thawed conditioned media was centrifuged at 3200g for 10min to pellet cell debris. Supernatant was collected and supplemented with sterile polyethylene glycol (PEG) 6000 buffer(*6, 77*) at 0.4ml buffer per 1ml conditioned media (to achieve a final PEG concentration of 8% w/v). Alternatively, the PEG-based miRCURY exosome isolation kit (Qiagen, 76743) was utilized with the same protocol and concentration for EV prep for miRNA sequencing.

PEG-media solutions were rotated overnight at 4°C then centrifuged for 30min at 3200g and 21°C to pellet EVs. Pellets were rinsed superficially with PBS without resuspension. Pelleted EVs were resuspended in an appropriate volume of resuspension buffer (Qiagen, 76743) to produce a 100x concentrated EV solution relative to the initial volume of conditioned media from which they were derived. Concentrated EVs were stored at -80°C. For isolation of EV total protein (western blotting) or miRNA (miRNA sequencing), EV pellets were resuspended directly in RIPA or Qiazol lysis buffer respectively, as previously described.(*6*)

### EV characterization

Aliquots of 100x concentrated EV solutions were diluted to 1x in PBS and nanoparticle tracking analysis (NTA) was used to quantify EV size distribution and concentration via a Nanosight NS300 (Malvern). TEM sample preparation and negative staining was performed as described previously, followed by imaging on a Talos L120C TEM (ThermoFisher Scientific) at 92,000x magnification.(*6*) Total protein quantification, western blotting, staining for CD63 or ALIX markers, and imaging was performed exactly as outlined previously, with the exception of MES SDS running buffer (Invitrogen) being used for the gel electrophoresis step.(*6*)

miRNA was isolated from lysed EV pellets using the miRNeasy Micro Kit (Qiagen, 217084) and sent to Novogene for DNase treatment, quantification, library preparation, sequencing, and read trimming according to a previously published protocol.(*78*) Biological replicates with fewer than 100 total aligned miRNA counts were considered to have failed sequencing and not included in further analyses (NT2, FGF1). The Qiagen RNA-seq Analysis Portal 3.0 (workflow version 1.2) was used to align trimmed reads to the human genome and to generate raw miRNA count files as described.(*78*) ComBat-seq R code was used to perform batch effect adjustment of raw miRNA count data based on RNA extraction cell culture batches according to a published protocol(*79*) and as outlined in the **Supplemental Code** file. EV-miRNA sequencing data was deposited in the GEO repository (accession: GSE300517).

For each sample, miRNA counts were normalized to total number of reads then converted to counts per million transcripts. PCA, differential expression analysis between each pair of sample groups, and heatmapping of miRNAs with a nominal p value<0.05 between any 2 groups was performed as outlined earlier, excluding the unstimulated ‘NT’ group from these analyses.

### Biowire ischemia-reperfusion injury induction and characterization

Epicardial Biowire tissues were subjected to IRI in vitro according to a previously established protocol.(*19, 35*) Briefly, 7 days after epicardial cell seeding (considered day 0), Biowire culture media was removed and replaced with 0.5 ml per well ischemic media solution consisting of 119 mM NaCl, 12 mM KCl, 1.2 mM NaH_2_PO_4_,1.3 mM MgSO_4_, 0.5 mM MgCl, 0.9 mM CaCl_2_, 20 mM sodium lactate, and 5 mM HEPES titrated to pH 6.4. Tissues were incubated in a hypoxic sub-chamber within a 37°C incubator supplied with 5% CO_2_ and 95% N_2_ and controlled by a ProOx P110 oxygen control unit (BioSpherix) set to the lowest possible setting (0.1% O_2_). Tissues were cultured in ischemic conditions for 6 hours. Ischemic media was then removed and replaced with 0.5 ml of reperfusion media per well consisting of RPMI 1640 supplemented with 1.4 mM CaCl_2_ and 2% (v/v) B27 without antioxidants (ThermoFisher, 10889038) titrated to pH 7.4. At this point, tissues were returned to normoxic culture in a 37°C 5% CO_2_ incubator. After 3 hours of reperfusion, media was changed back to the regular 1ml of I3M culture media and tissues were cultured at 37°C and 5% CO_2_ until the experimental endpoint for RNA extraction or immunofluorescence imaging one week after IRI. Control uninjured tissues were cultured in parallel for the same time period without IRI exposure for comparative endpoint analysis (ie. 15 days after initial CM/CF cell seeding).

### Epicardial-EV supplementation of Biowire tissues

Epi-EVs from each stimulation conditioned were isolated and quantified by NTA as described above. For the preliminary Epi-EV functional screening experiment, doses of 4x10^8^ concentrated NT-EVs, SB-EVs, TGF-β-EVs, or FGF-EVs were added directly to the I3M culture media of separate Epicardial Biowires immediately upon completion of the full IRI protocol to achieve an EV dose as close in magnitude as possible to previously published studies while balancing variable EV yield from each epicardial-derived cell source.(*78, 80, 81*) 0.5 ml of additional fresh I3M media was added to Biowire tissue four days after IRI (including non-EV supplemented controls for consistency).

For the follow-up, targeted SB-EV supplementation experiment, SB-EVs were dosed at 1.25x10^9^ EVs per tissue based on previously published protocols.(*80, 81*) A half dose of SB-EVs (6.25x10^8^ EVs/tissue) was added to each Epicardial Biowire tissue well during ischemia and again during reperfusion (to account for the reduced media volume of 0.5ml during each step, maintaining a consistent EV concentration). When tissues were returned to the regular 1ml of I3M after IRI, a full dose of SB-EVs (1.25x10^9^ EVs per tissue) was added directly to the Biowire culture media. Media was removed on day 3 and replaced with fresh I3M (1ml + 1.25x10^9^ SB-EVs per well). This media change was also performed at the same time for non-EV supplemented control tissues.

### Visualization of EV uptake in Biowire tissues

150μl total of pooled NT- and FGF-EVs (100x concentrated) were mixed at a 1:3 ratio by volume with Cell Tracker CM-DiI (Invitrogen C7001; 1 mg/ml in DMSO, diluted to 5μg/ml with PBS) and incubated for 15 minutes in the dark at room temperature.

Stained Epi-EVs were then added to an Amicon Ultra-0.5 ml 100kDa filter unit (MilliporeSigma UFC510024) and centrifuged for 5 min at 14 000 g. Two PBS washes of EVs were performed by adding 500μl PBS to the filter unit, mixing gently, then centrifuging for 5 min at 14 000 g. The filter unit was then flipped upside-down into a fresh collector tube and centrifuged for 2 min at 1000 g to collect washed and stained Epi-EVs.

Biowire tissues (CM + CF) were seeded according to the protocol described earlier, with the exception of BJ1D iPSC-derived CMs being utilized for this supporting experiment instead of iCell CMs. BJ1D iPS-CMs were differentiated as outlined in a previously published protocol.(*19*)

Stained EVs were added to 500μl Biowire tissue culture media 3 days after initial seeding and incubated overnight. The following day, EV-supplemented and non-EV supplemented tissues were both washed with PBS, gently removed from wires using a P1000 pipette tip, and pipetted into separate 1.7ml microcentrifuge tubes. PBS was gently removed and 4% PFA was added to the tubes for tissue fixation at room temperature in the dark for 1 hour. Fixed tissues were rinsed with PBS then stained with DAPI (1:500, 10 min), washed with PBS three times for 5 min each, then placed in a confocal dish and compressed gently with a glass coverslip. Tissues were imaged on a Zeiss LSM710 two-photon microscope in the DAPI and AF568 channels, using the exact same imaging and display settings for tissues with and without stained Epi-EVs.

### Tissue functional assessment and GFP cell tracking

Functional assessment of Epicardial Biowire tissue contractile properties was performed before IRI as well as 12 hours, 3 days, and 7 days after IRI as outlined in detail previously.(*19, 21, 67*) Briefly, a microscale mechanical tester was used to define the mechanical properties of polymer microwires before tissue seeding. During assessments, Biowire plate electrodes were connected to an electrical stimulator under a fluorescent microscope. Tissue dimensions were measured in the brightfield channel. Excitation threshold voltage (ETV) was measured by setting electrical stimulation to 1V and 1 Hz and increasing the voltage 0.1V at a time until tissues beat synchronously in time with applied electrical pulses (at which point, the voltage was recorded as ETV).

Tissues that exceeded 8V at 1Hz and still did not beat synchronously with applied pulses were considered ‘not catching’. Maximum capture rate (MCR) was assessed by setting the stimulation voltage at 2x the ETV (up to a maximum of 8V) and slowly ramping up frequency of stimulation from 1Hz, increasing by 0.1Hz at a time until tissues no longer beat synchronously with applied pulses (at which point, MCR was recorded as the frequency in Hz just before synchronicity was lost). The remainder of contractile properties were assessed at 1Hz and 2x the ETV. Videos of wire deflection under this stimulation protocol were recorded under the Texas red fluorescent channel and custom MATLAB codes were utilized to analyze wire deflection and compute passive tension, contractile force (active tension) and contraction/relaxation slopes as described, based on measured polymer wire mechanical properties.(*19, 67*) Functional properties of each tissue were normalized to the individual tissue’s cross sectional area as well as its ‘before IRI’ value for relative comparisons between tissues/groups.

At the same timepoints as the functional analyses described above, tissues were imaged in the brightfield and FITC channels of the fluorescent microscope for GFP+ epicardial-derived cell tracking within tissues as outlined previously.(*19*) Briefly, FITC channel images of Epicardial Biowires were processed in Fiji for background subtraction and thresholding. Plot profile was performed at three locations across the width of each Biowire, with pixel values converted to binary and binned at 10% intervals to determine GFP+ pixel distribution across tissue width.(*19*)

### Tissue immunostaining and microscopy

Representative Epicardial Biowire tissues not lysed for RNA sequencing were harvested on day 7 after IRI (using the same procedure to detach tissues from wires as described before the RNA extraction protocol outlined above) and fixed in 4% PFA at 4°C overnight followed by several PBS washes. Tissues were then prepared for immunostaining and fluorescent imaging according to a published protocol.(*19*) Briefly, tissues were cut in half with a razor blade and blocked for 1 hour at room temperature with gentle rocking in 10% NGS + 0.1% TritonX-100. Primary staining was performed in 100 μL of the same blocking buffer at 4°C overnight with the addition of anti-cardiac troponin T (Invitrogen, MA5-12960, 1:100, mouse monoclonal) or anti-vimentin (Abcam, ab8978, 5µg/ml, mouse monoclonal) to separate tissue halves. Primary antibody solution was removed and tissues were stored in fresh PBS at 4°C overnight.

Secondary antibody staining was then performed in 100 μL of blocking buffer at 4°C overnight with the addition of goat anti-mouse AF647-conjugated secondary (Invitrogen, A21240, 1:200). Another overnight PBS wash was performed as before, tissues were stained with DAPI (1:1000), washed with PBS, then imaging was performed on a Nikon A1R confocal microscope.

### miRNA target prediction

The top 20 most abundant SB-EV miRNAs were searched in the miRDB database(*39*) (https://mirdb.org/mining.html; accessed January 2025) and predicted mRNA targets with a target score greater than or equal to 60 (the default value for the target mining feature) were filtered by the list of significantly downregulated genes in the ‘Epi + IRI + SB-EVs’ tissue RNA sequencing group. The same process of top 20 SB-EV miRNA searching and downregulated tissue mRNA target filtering was performed on the following databases: the DIANA-microT 2023 webserver(*40*) (https://dianalab.e-ce.uth.gr/microt_webserver/#/interactions; accessed January 2025, miRBase v22.1 database) with an interaction score ≥0.70; TargetScan(*41, 82*) (https://www.targetscan.org/vert_80/; accessed January 2025, Release 8.0 September 2021) with a weighted context++ score <-0.1; miRTarBase(*42*) 2025 (https://awi.cuhk.edu.cn/~miRTarBase/miRTarBase_2025/php/index.php; accessed January 2025); and TarBase(*43*) (https://dianalab.e-ce.uth.gr/tarbasev9/interactions; accessed January 2025, v9.0). For each given miRNA-mRNA interaction pair, the number of predictive database hits (within the listed cut-off scores) and experimental database hits was added together then filtered to highlight interactions with 3 or more total hits between the 5 databases queried. The total number of database hits for any given top 20 SB-EV miRNA and for any downregulated Epi + IRI + SB-EVs mRNA were also separately added together to rank and plot the most predicted miRNAs and genes relevant to SB-EV supplementation of Epicardial Biowire tissues.

The Broad Institute GSEA(*32*) mac app (https://www.gsea-msigdb.org/gsea/downloads.jsp; v4.3.2, October 2022) was used for GSEA of miRNA targets in Epi + IRI vs. Epi + IRI +SB-EVs mRNA sequencing data (pre-ranked GSEA, using previous ranked gene list from earlier mRNA sequencing analysis; applying c3.mir.mirdb.v2024.1.Hs.symbols.gmt MSigDB(*83*) collection as a reference database). Resulting enriched miRNA target gene sets (FDR<0.25) were filtered by those that appeared in the top 20 SB-EV miRNA list.

### Data and Statistical Analyses

Statistical analyses and plotting were performed using GraphPad Prism (v10.5.0) unless indicated otherwise. Data are presented as mean ± standard deviation (SD) with sample sizes indicated in figure captions. For Epi-EV NTA and epicardial cell immunofluorescence analyses, the Shapiro-Wilk test was used to check normality, and data that passed was analyzed via one-way ANOVA with a post-hoc Tukey test for multiple comparisons as indicated in figure captions. Where data did not pass the Shapiro-Wilk normality test, the Kruskal Wallis nonparametric test was used followed Dunn’s multiple comparisons test. For Biowire functional tissue analyses, repeated measures two-way ANOVA or mixed effects model (where appropriate) analyses were performed, assuming a normal distribution of residuals with a Geisser-Greenhouse correction (ie. no assumption of sphericity) followed by post-hoc Tukey’s multiple comparison tests (individual variances computed for each comparison). P<0.05 or FDR-adjusted P<0.05 (where appropriate) was considered significant, except for GSEA where the threshold was set to FDR<0.25 to align with the GSEA-specific recommendation.(*32*)

