## Supplemental Figures for "Epicardial extracellular vesicles modulate gene expression following ischemia-reperfusion injury in heart-on-a-chip"

**Supplemental Information**

Supplemental Data Tables

Supplemental data tables can be found in the attached file.

Supplemental Figures

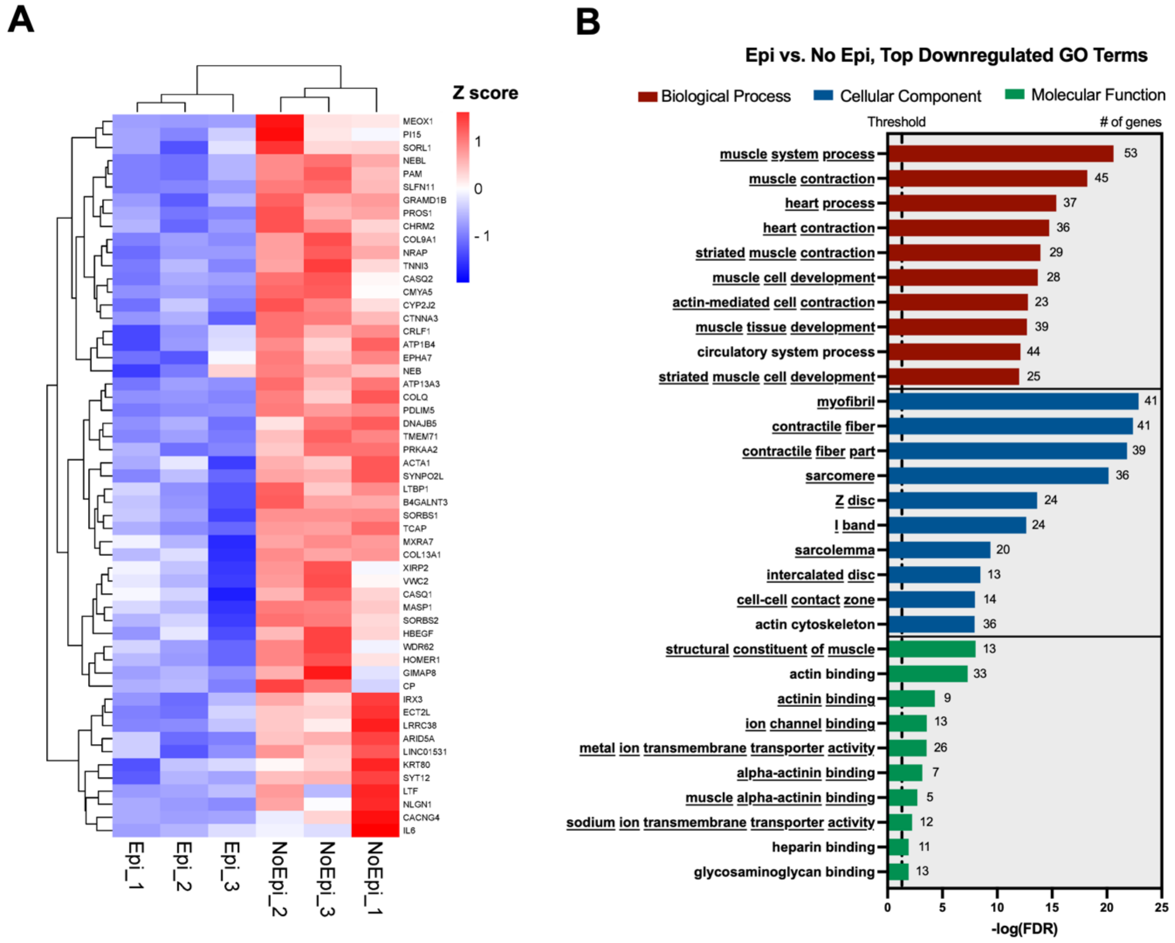


**Supplemental Figure S1. Downregulated genes and GO terms highlighted cardiac-specific transcripts and processes in heart-on-a-chip containing epicardial cells** which, by nature, included a smaller proportion of cardiomyocytes. (A) Heatmap of all significantly downregulated genes in Epi vs. No Epi tissues. (B) Top 10 downregulated GO terms in Epi vs. No Epi tissues. Underlined text indicates that a given GO term is uniquely significant to downregulated terms (ie. enriched in No Epi)

and is not enriched in Epi tissues. For (A) and (B): n=3 for both groups; FDR<0.05 considered significant.

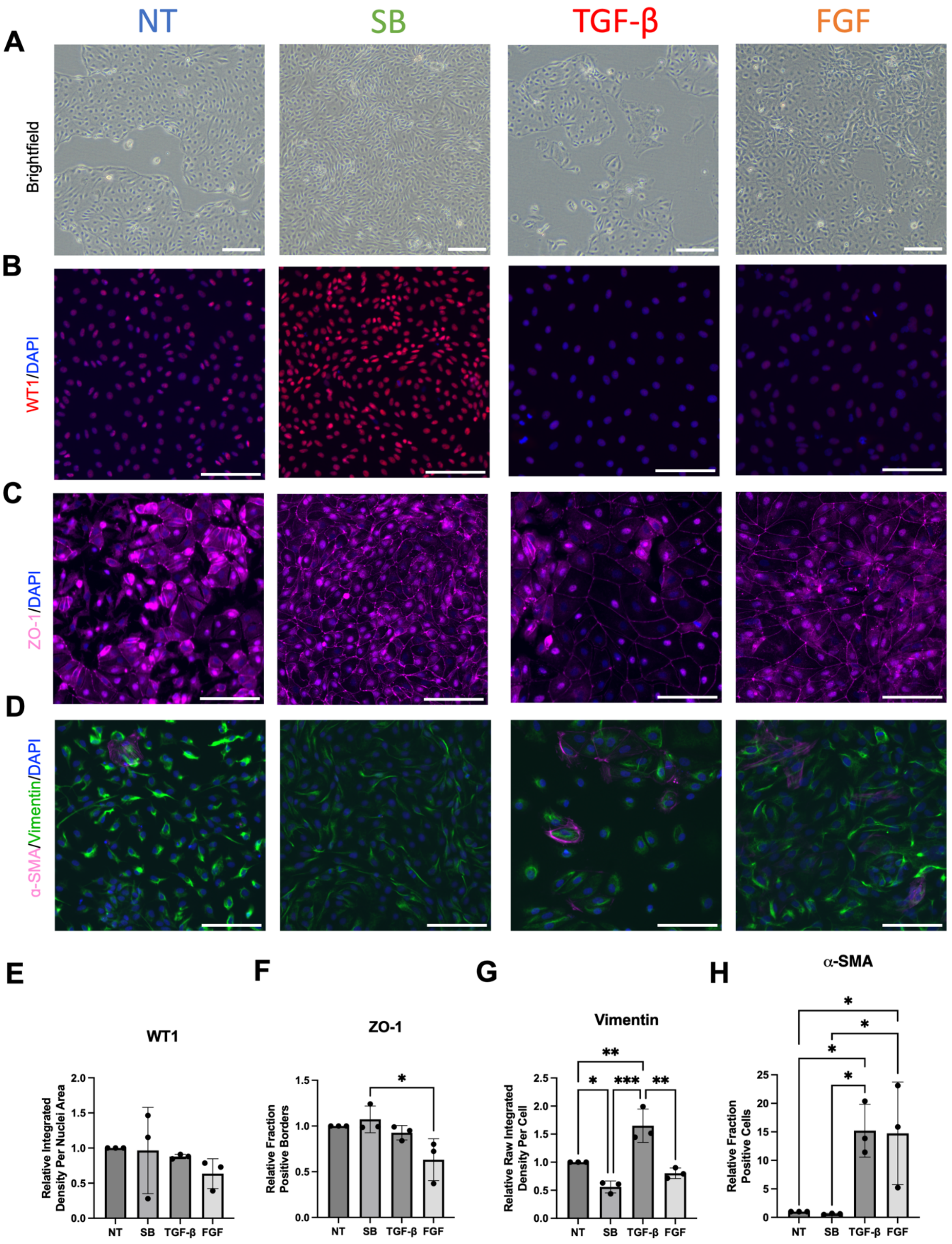


**Supplemental Figure S2. Cell monolayers after differential stimulation to create three different linages of epicardial-derived cells**. No treatment control cells ('NT');

epicardial cells (stimulated with SB-431542, a TGF-β inhibitor, to complete epicardial differentiation; ‘SB’); EMT-stimulated with transforming growth factor-β; ‘TGF-β’; and EMT-stimulated with basic fibroblast growth factor; ‘FGF’. (A) Brightfield images of epicardial derived cells fixed on day 8 (NT, SB) or day 9 (TGF-β, FGF) of culture. Scale bars: 200μm. (B-D) Immunofluorescence (IF) imaging of epicardial derived cells fixed on day 8 (NT, SB) or day 9 (TGF-β, FGF) of culture. Scale bars: 100μm. Cells were stained to highlight nuclei (DAPI, blue), in addition to: (B) Wilms tumor protein (WT1, red), (C) zonula occludens-1 (ZO-1, pink), (D) vimentin (green), and α-smooth muscle actin (α-SMA, pink). (E-H) Quantification of IF images: (E) integrated density of WT1 signal per nuclei area relative to NT control cells from a given batch; (F) fraction of cells with ZO-1 positive borders relative to NT control cells from a given batch; (G) raw integrated density of vimentin signal per cell relative to NT control cells from a given batch; (H) fraction of α-SMA positive cells relative to NT control cells from a given batch. For (E): Kruskal Wallis test (nonparametric) with post-hoc Dunn’s multiple comparisons test. For (F)-(H): One-way ANOVA with post-hoc Tukey test; * P<0.05, **P<0.01, ***P<0.001. Data presented as mean ± standard deviation (SD).

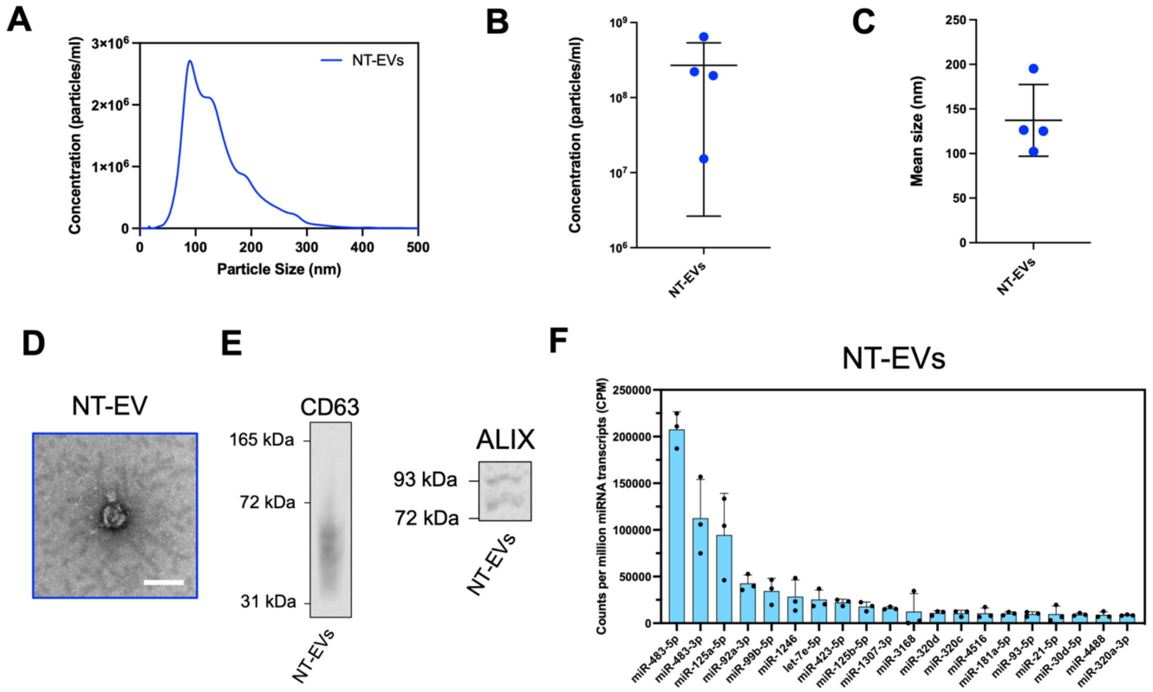

**Supplemental Figure S3. Characterization of extracellular vesicles isolated from unstimulated pro-epicardial cell controls (NT-EVs).** (A-C) NTA results for (A) size distribution, (B) concentration, and (C) mean size of NT-EVs. N=4. (D) TEM image of NT-EVs. Scale bar: 100nm. (E) Western blot detection of CD63 and ALIX in NT-EV isolate. (F) Top 20 most abundant miRNAs detected in miRNA sequencing of NT-EVs. N=3. Data presented as mean ± standard deviation (SD).

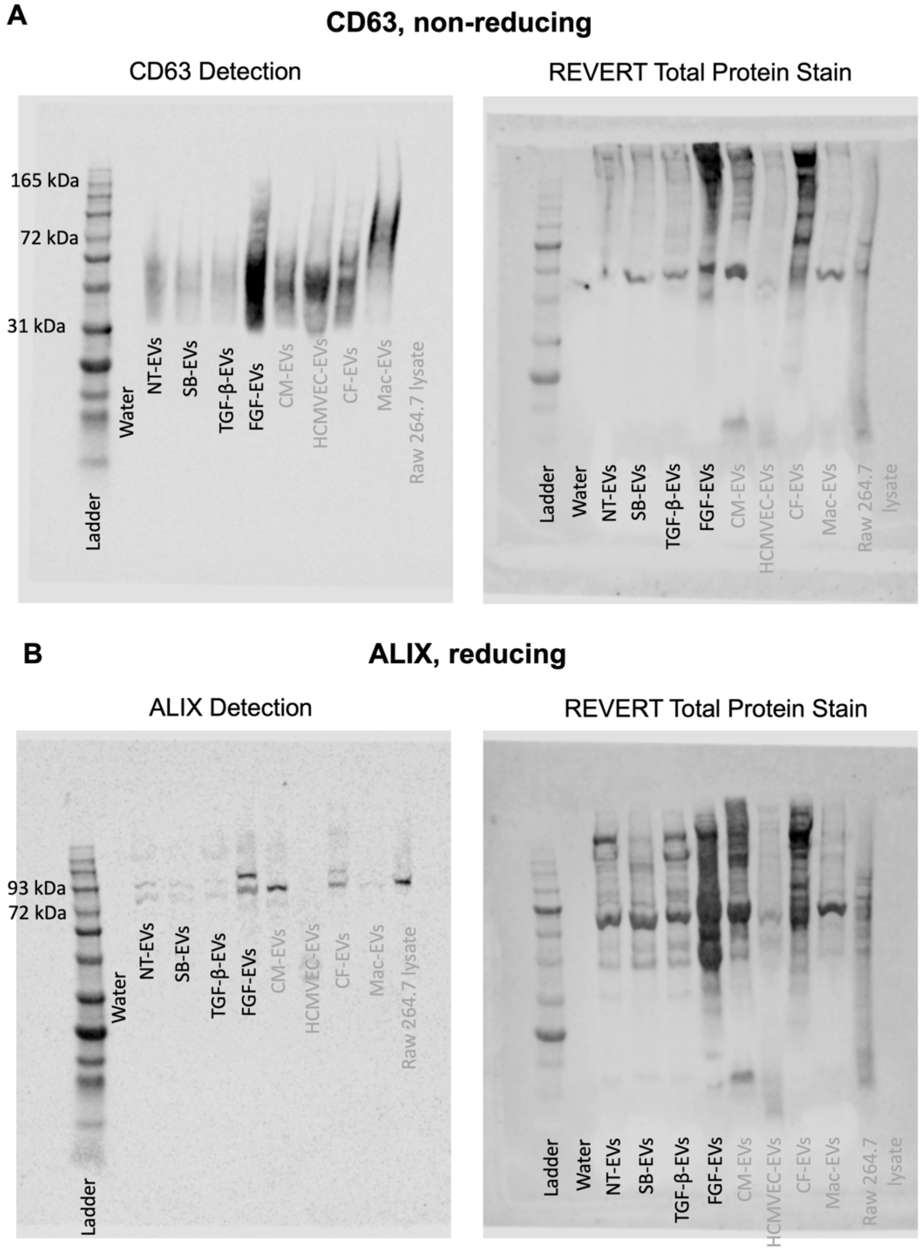

**Supplemental Figure S4. Full western blot images depicting Epi-EV protein marker signal as well as total protein stain detection** for (A) CD63 (blot run in non-reducing conditions) and (B) ALIX (blot run in reducing conditions). Lanes labelled with grey letters are outside the scope of this manuscript.

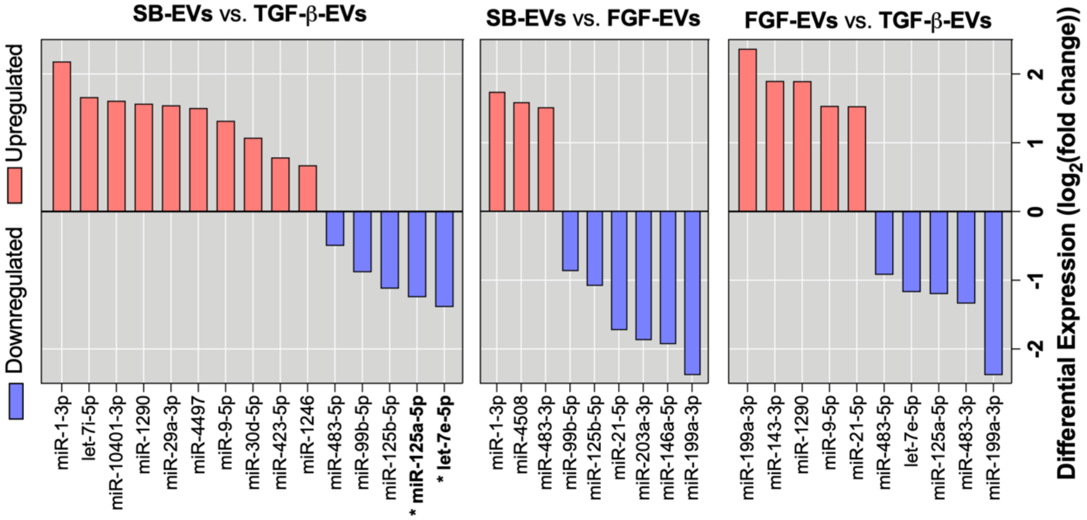


**Supplemental Figure S5. Detailed results from differential expression analysis of epicardial-EV miRNA sequencing** showing all miRNAs identified to have a nominal p-value<0.05 from pairwise comparisons of each of the three EV types. * indicates FDR<0.05.

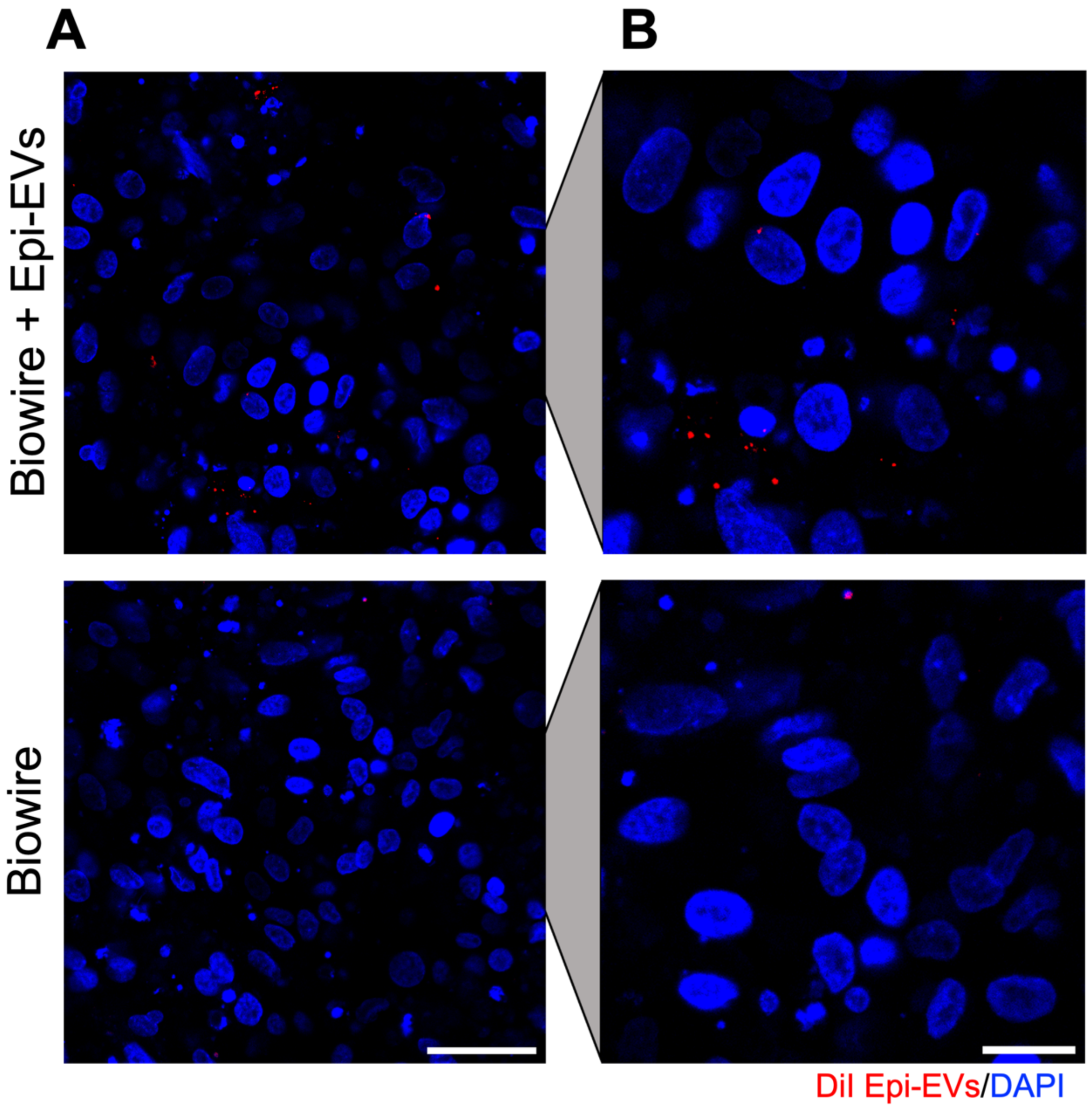


**Supplemental Figure S6**. **Biowire heart-on-a-chip uptakes extracellular vesicles.** (A,B) Representative images showing uptake of DiI-stained Epi-EVs (red, incubated overnight) in Biowire tissue compared to tissue without EV supplementation. Cell nuclei were stained with DAPI (blue). (A) scale bar: 50μm; (B) scale bar: 20μm.

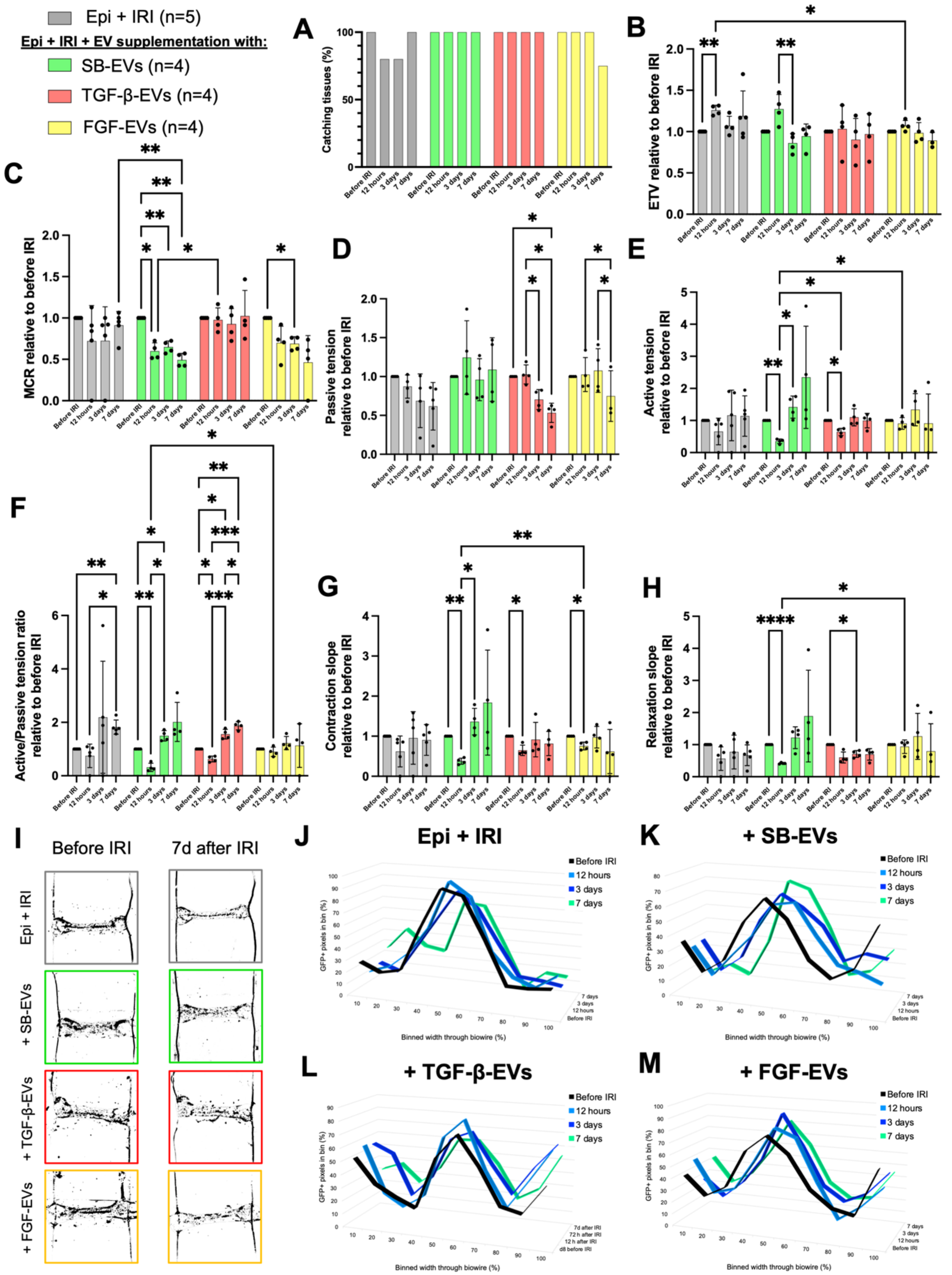


**Supplemental Figure S7. Contractility and epicardial cell migration in hearts-on-a-chip supplemented with SB-EVs, TGF-β-EVs, or FGF-EVs post-IRI.** (A-H) Characterization of (A) percentage of Biowire tissues 'catching' electrical stimulation paced at 1Hz, (B) excitation threshold voltage (ETV), (C) maximum capture rate (MCR), (D) passive tension, (E) active tension, (F) active/passive tension ratio, (G) contraction slope, and (H) relaxation slope of each engineered Biowire tissue at 12 hours, 3 days, and 7 days post-IRI. N= 5 for Epi + IRI(*19*); n=4 for Epi + IRI + SB-EVs, Epi + IRI + TGF-β-EVs, and Epi + IRI + FGF-EVs. (I) Representative images of Biowire tissues before and 7 days after IRI/EV supplementation illustrating how thresholding of GFP+ epicardial cells was used to perform epicardial cell tracking through the width of Biowire tissues over time and with epicardial derived cell-EV supplementation. (J-M) Averaged distribution of GFP+ epicardial cells throughout Biowire tissue width for (J) Epi + IRI injured control tissues(*19*) and injured tissues supplemented with (K) SB-EVs, (L) TGF-β-EVs, or (M) FGF-EVs before IRI and 12 hours, 3 days, and 7 days after IRI. N=4 for all groups. (B)-(H): properties are shown relative to before IRI for each tissue and set to zero for tissues that did not catch electrical stimulation at 1Hz (except for (B) and (D), where non-catching tissues were discarded from analyses). Repeated measures two-way ANOVA (C, E-H) or mixed effects model (B,D) with Geisser-Greenhouse correction; post-hoc Tukey's multiple comparison test (individual variances computed for each comparison). *P<0.05, **P<0.01, ***P<0.001, ****P<0.0001. Data presented as mean ± standard deviation (SD).

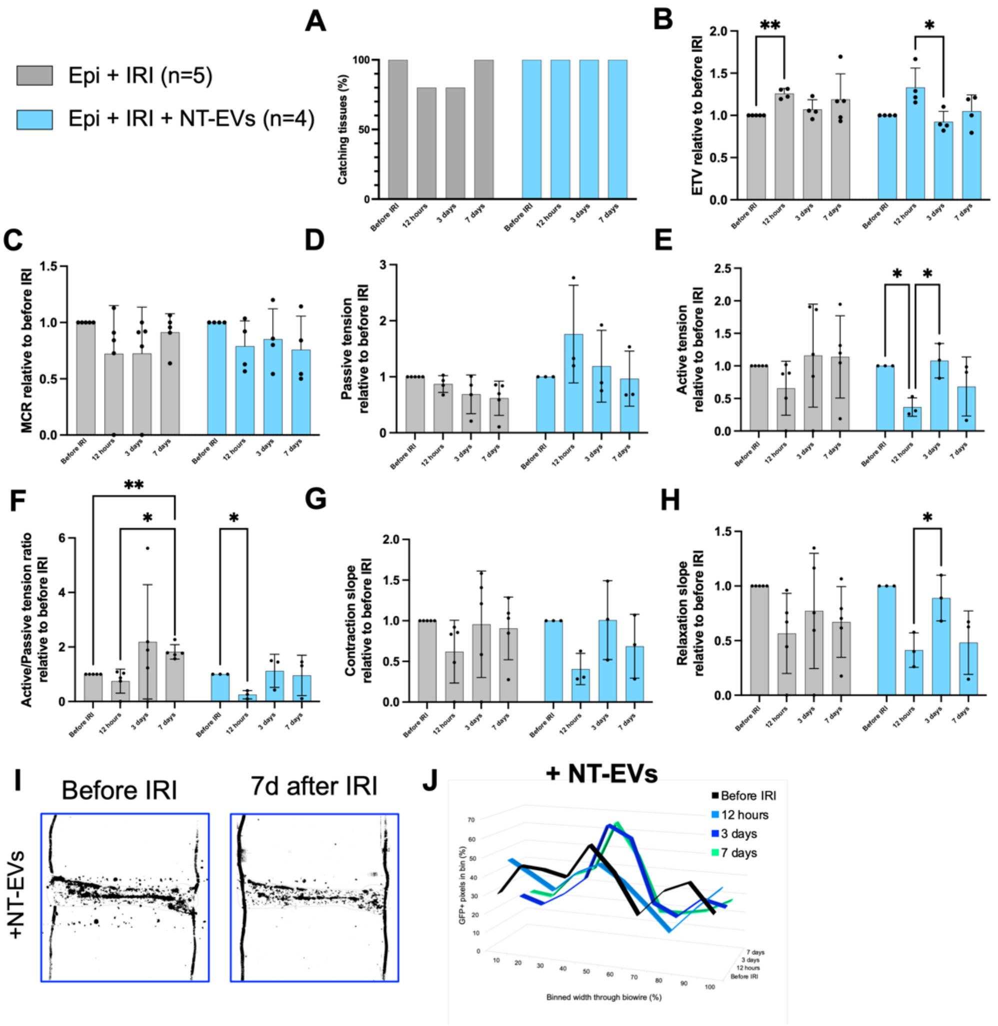


**Supplemental Figure S8. Contractility and epicardial cell migration in hearts-on-a-chip supplemented with NT-EVs.** (A-H) Characterization of (A) percentage of Biowire tissues ‘catching’ electrical stimulation paced at 1Hz, (B) excitation threshold voltage (ETV), (C) maximum capture rate (MCR), (D) passive tension, (E) active tension, (F) active/passive tension ratio, (G) contraction slope, and (H) relaxation slope of each engineered Biowire tissue at 12 hours, 3 days, and 7 days post-IRI. N= 5 for Epi + IRI(*19*); n=4 for Epi + IRI + NT-EVs. (I) Representative images of Biowire tissues before and 7 days after IRI/EV supplementation illustrating how thresholding of GFP+ epicardial cells was used to perform epicardial cell tracking through the width of Biowire tissues over time and with NT-EV supplementation. (J) Averaged distribution of GFP+ epicardial cells throughout Biowire tissue width for injured tissues supplemented with NT-EVs before IRI and 12 hours, 3 days, and 7 days after IRI. N=4. (B)-(H): properties are shown relative to before IRI for each tissue and set to zero for tissues that did not catch electrical stimulation at 1Hz (except for (B) and (D), where non-catching tissues were discarded from analyses). Repeated measures two-way ANOVA (C, E-H) or mixed effects model (B,D) with Geisser-Greenhouse correction; post-hoc Tukey's multiple comparison test (individual variances computed for each comparison). *P<0.05, **P<0.01. Data presented as mean ± standard deviation (SD).

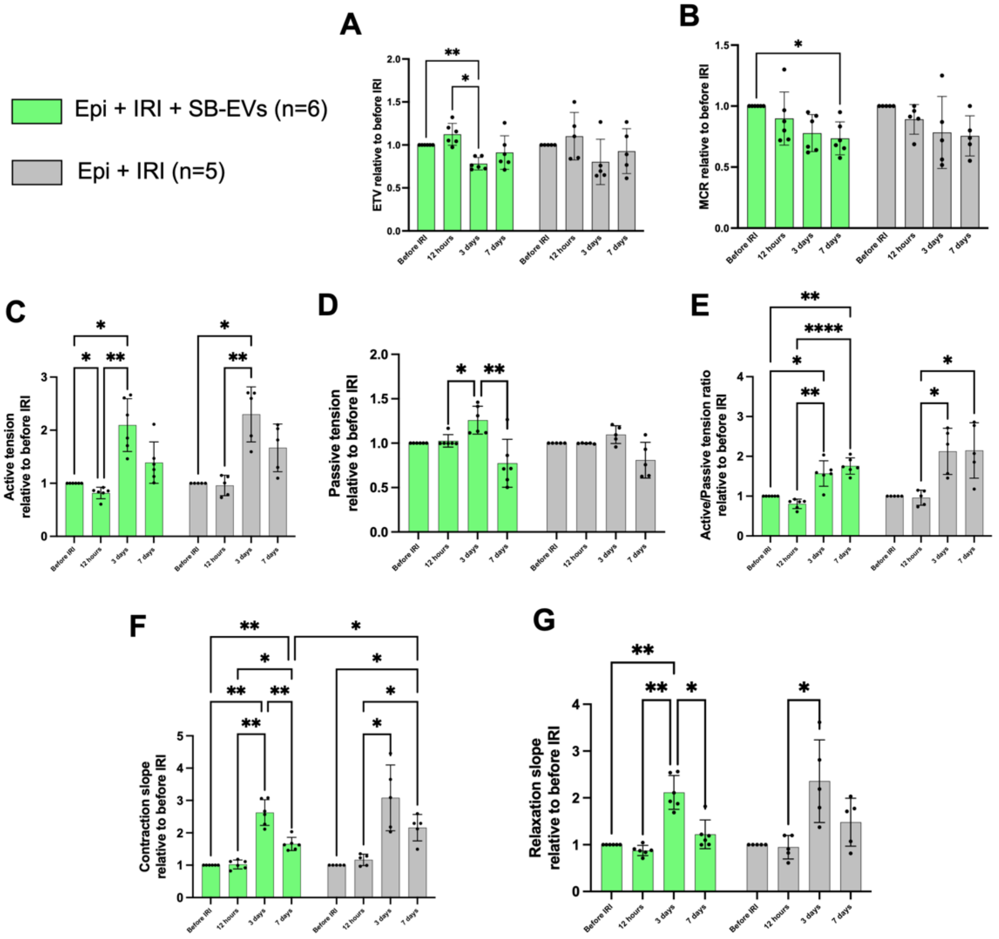


**Supplemental Figure S9. Targeted functional analyses of SB-EV supplemented hearts-on-a-chip post IRI did not reveal significant changes in tissue contractility or excitability when compared to IRI controls without EV supplementation**. (A-G) Characterization of (A) excitation threshold voltage (ETV), (B) maximum capture rate (MCR), (C) active tension, (D) passive tension, (E) active/passive tension ratio, (F) contraction slope, and (G) relaxation slope of each engineered Biowire tissue at 12 hours, 3 days, and 7 days post-IRI relative to before IRI. N= 6 for Epi + IRI + SB-EVs, n=5 for Epi + IRI. Repeated measures two-way ANOVA with Geisser-Greenhouse correction; post-hoc Tukey's multiple comparison test (individual variances computed for each comparison). *P<0.05, **P<0.01, ***P<0.001, ****P<0.0001. Data presented as mean ± standard deviation (SD).

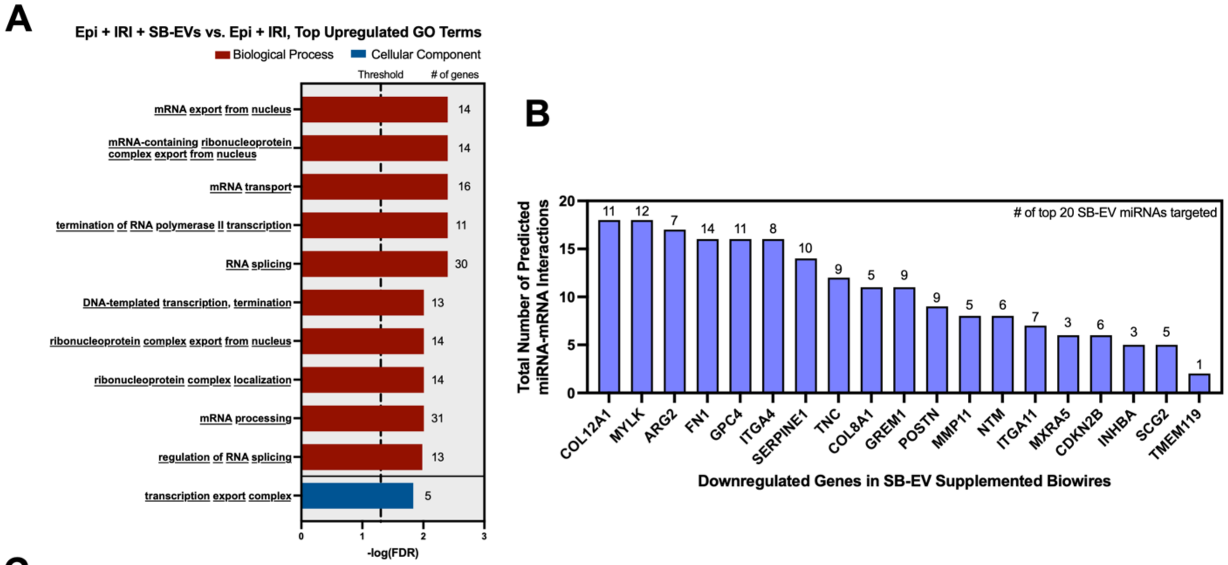


C

| miRNA | mRNA | Predictive Databases | | | | Experimental Databases | | # of Predictive Database Hits | # of Experimental Database Hits | Total Number of Database Hits |
|---|---|---|---|---|---|---|---|---|---|---|
| | | miRDB | microT | TargetScan | | miRTarBase | TarBase | | | |
| | | Target Score (>60) | Interaction Score (>0.70) | Conservation Status | Weighted Context++ Score (<-0.1) | Total # of Experiments | Total # of Experiments | | | |
| let-7b-5p | ARG2 | 81 | 0.88 | Conserved | -0.4 | - | 4 | 3 | 1 | **4** |
| let-7e-5p | ARG2 | 81 | 0.84 | Conserved | -0.4 | - | 4 | 3 | 1 | **4** |
| miR-16-5p | COL12A1 | 89 | 0.81 | Conserved | -0.22 | - | 8 | 3 | 1 | **4** |
| miR-16-5p | MYLK | 96 | 0.83 | Conserved | -0.42 | - | 13 | 3 | 1 | **4** |
| miR-9-5p | COL12A1 | 73 | 0.80 | - | - | 1 | 2 | 2 | 2 | **4** |
| miR-9-5p | TNC | 98 | 1.00 | Conserved | -0.27 | - | 1 | 3 | 1 | **4** |
| let-7b-5p | COL8A1 | - | - | Poorly Conserved | -0.13 | 2 | 2 | 1 | 2 | **3** |
| let-7e-5p | COL8A1 | - | - | Poorly Conserved | -0.13 | 2 | 3 | 1 | 2 | **3** |
| miR-10a-5p | ARG2 | 68 | 0.79 | Poorly Conserved | -0.55 | - | - | 2 | 0 | **3** |
| miR-10a-5p | SERPINE1 | - | 0.80 | - | - | 3 | 2 | 1 | 2 | **3** |
| miR-21-5p | GPC4 | 66 | 0.74 | Conserved | -0.14 | - | - | 3 | 0 | **3** |
| miR-30d-5p | MXRA5 | 86 | 0.95 | - | - | - | 1 | 2 | 1 | **3** |
| miR-30d-5p | COL8A1 | 77 | - | Poorly Conserved | -0.19 | - | 3 | 2 | 1 | **3** |
| miR-30d-5p | ITGA4 | - | 0.96 | Conserved | -0.24 | - | 1 | 2 | 1 | **3** |
| miR-30d-5p | NTM | - | 0.91 | Poorly Conserved | -0.15 | - | 3 | 2 | 1 | **3** |
| miR-30d-5p | SERPINE1 | - | - | Conserved | -0.14 | 5 | 13 | 1 | 2 | **3** |
| miR-320c | ITGA4 | - | 0.83 | Poorly Conserved | -0.14 | - | 1 | 2 | 1 | **3** |
| miR-9-5p | ITGA4 | 87 | 1.00 | - | - | - | 1 | 2 | 1 | **3** |
| miR-9-5p | MYLK | - | - | Conserved | -0.12 | 1 | 1 | 1 | 2 | **3** |

**Supplemental Figure S10. Predicted miRNA-mRNA interactions in IRI injured hearts-on-a-chip supplemented with SB-EVs.** (A) Enriched GO terms for genes upregulated in Epi + IRI + SB-EV tissues compared to Epi + IRI controls. N=3 for both groups; FDR<0.05 considered significant. Underlined text indicates that a given term is uniquely significant to upregulated GO terms and is not significantly enriched in GO analysis of downregulated DEGs. (B) Plot of Epi + IRI + SB-EV downregulated genes with the greatest total number of predicted miRNA-mRNA interactions for all abundant and enriched SB-EV miRNAs from all five predictive/experimental miRNA databases presented in Table 1. Numbers above each bar indicate the number of miRNAs with at least one predicted/experimental miRNA-mRNA interaction with a given gene (out of 26 total key SB-EV miRNAs searched). (C) Extended list of correlated miRNA-mRNA interactions between abundant and enriched SB-EV miRNAs and downregulated genes in Epi + IRI + SB-EV tissues with at least 3 predictive or experimental database hits, depicting results from specific databases, scores, and cut-offs applied where appropriate.
