## Supplemental Code for "Epicardial extracellular vesicles modulate gene expression following ischemia-reperfusion injury in heart-on-a-chip": ComBat_Seq_KW_052923.html

Batch Effect Adjustment in R Using Combatseq


### Batch Effect Adjustment in R Using Combatseq

Citation: Zhang Y, Parmigiani G, Johnson WE. ComBat-seq: Batch effect
adjustment for RNA-seq count data. NAR Genomics Bioinforma.
2020;2(3):1-10. doi:10.1093/nargab/lqaa078

Performed By: Karl Wagner

Date: May 29, 2023

Description: Batch effect adjustment of Epi-EV miRNA seq raw counts
according to RNA extraction and epicardial culture batches.

### RNA extraction batch effect adjustment

Load necessary packages

```
library("sva")
```

```
## Loading required package: mgcv
```

```
## Loading required package: nlme
```

```
## This is mgcv 1.9-3. For overview type 'help("mgcv-package")'.
```

```
## Loading required package: genefilter
```

```
## Loading required package: BiocParallel
```

```
library("ggplot2")
```

Load the raw miRNA count data

```
uncorrected_data = read.table("Uncorrected_rawcounts_all4.txt", header=TRUE, sep="\t", as.is=c(1))

#First column - keep as characters
```

```
names(uncorrected_data) = c("Name", "FGF2", "FGF3", "FGF4", "NT1", "NT3", "NT4", "TGFb1", "TGFb2", "TGFb3", "TGFb4", "SB1", "SB2", "SB3", "SB4")
sample_names = names(uncorrected_data)[2:length(names(uncorrected_data))]
```

Review data structure

```
head(uncorrected_data)
```

```
dim(uncorrected_data)
```

```
## [1] 2632   15
```

Define samples, conditions (groups), and batches

```
conditions = c("FGF", "FGF", "FGF", "NT", "NT", "NT", "TGFb", "TGFb", "TGFb", "TGFb", "SB", "SB", "SB", "SB")

#Define RNA extraction batches
RNA_batch = c(1, 2, 2, 1, 2, 2, 1, 1, 2, 2, 1, 1, 2, 2)

#Define epicardial cell culture batches
replicates = c(2, 3, 4, 1, 3, 4, 1, 2, 3, 4, 1, 2, 3, 4)
```

Transform the format of groups and batches from names to numbers

```
#in the command below "sapply" is used to apply the "switch" command to each element and convert names to numbers as we define
#for conditions:
groups = sapply(as.character(conditions), switch, "FGF" = 1, "NT" = 2, "TGFb" = 3, "SB" = 4, USE.NAMES = F)
```

Run Combatseq for RNA extraction batch

```
corrected_data = ComBat_seq(counts = as.matrix(uncorrected_data[,sample_names]), batch = RNA_batch, group = groups)
```

```
## Found 2 batches
## Using full model in ComBat-seq.
## Adjusting for 3 covariate(s) or covariate level(s)
## Estimating dispersions
## Fitting the GLM model
## Shrinkage off - using GLM estimates for parameters
## Adjusting the data
```

Join the gene names onto the now corrected counts from ComBat\_seq

```
corrected_data = cbind(uncorrected_data[,c("Name")], corrected_data)
```

```
#compare dimensions of corrected and uncorrected data sets
dim(uncorrected_data)
```

```
## [1] 2632   15
```

```
dim(corrected_data)
```

```
## [1] 2632   15
```

```
#visually compare values of corrected and uncorrected data sets
head(uncorrected_data)
```

```
head(corrected_data)
```

```
##                    FGF2 FGF3 FGF4 NT1  NT3  NT4  TGFb1 TGFb2 TGFb3 TGFb4 SB1
## [1,] "let-7a-2-3p" "0"  "0"  "0"  "0"  "0"  "0"  "0"   "0"   "0"   "0"   "0"
## [2,] "let-7a-3p"   "0"  "0"  "0"  "0"  "0"  "0"  "0"   "0"   "0"   "0"   "0"
## [3,] "let-7a-5p"   "1"  "17" "7"  "45" "47" "10" "8"   "3"   "45"  "28"  "3"
## [4,] "let-7b-3p"   "0"  "0"  "0"  "0"  "0"  "0"  "0"   "0"   "0"   "0"   "0"
## [5,] "let-7b-5p"   "2"  "84" "5"  "4"  "19" "8"  "4"   "5"   "23"  "23"  "5"
## [6,] "let-7c-3p"   "0"  "0"  "0"  "0"  "0"  "0"  "0"   "0"   "0"   "0"   "0"
##      SB2 SB3  SB4 
## [1,] "0" "0"  "0" 
## [2,] "0" "0"  "0" 
## [3,] "4" "10" "5" 
## [4,] "0" "0"  "0" 
## [5,] "6" "14" "21"
## [6,] "0" "0"  "0"
```

Convert to dataframe

```
# Convert the character matrix to a data frame
corr_dataframe <- data.frame(corrected_data, stringsAsFactors = FALSE)
```

Write to excel file

```
library("writexl")
# Set the filename for the Excel file
filename <- paste0(getwd(), "/BatchCorrCounts.xlsx")
# Save the data frame to an Excel file
write_xlsx(corr_dataframe, path = filename)
```

### Cell culture batch effect adjustment

Load the RNA extraction batch effect adjusted count data

```
culture_uncorrected = read.table("RNABatchCorr_Counts_all4.txt", header=TRUE, sep="\t", as.is=c(1))

#First column - keep as characters
```

```
names(culture_uncorrected) = c("Name", "FGF2", "FGF3", "FGF4", "NT1", "NT3", "NT4", "TGFb1", "TGFb2", "TGFb3", "TGFb4", "SB1", "SB2", "SB3", "SB4")
culture_sample_names = names(culture_uncorrected)[2:length(names(culture_uncorrected))]
```

Review data structure

```
head(culture_uncorrected)
```

```
dim(culture_uncorrected)
```

```
## [1] 2632   15
```

Define samples, conditions, and batches as before

```
#same as before - don't need to re-define
#conditions = c("FGF", "FGF", "FGF", "NT", "NT", "NT", "TGFb", "TGFb", "TGFb", "TGFb", "SB", "SB", "SB", "SB")
#RNA_batch = c(1, 2, 2, 1, 2, 2, 1, 1, 2, 2, 1, 1, 2, 2)

#This second step will focus on culture batches as defined below:
culture_batch = c(2, 3, 4, 1, 3, 4, 1, 2, 3, 4, 1, 2, 3, 4)
```

Transform the format of groups and batches from names to numbers

```
#same as before
#in the command below "sapply" is used to apply the "switch" command to each element and convert names to numbers as we define
#for conditions:
#groups = sapply(as.character(conditions), switch, "FGF" = 1, "NT" = 2, "TGFb" = 3, "SB" = 4, USE.NAMES = F)
```

Run Combatseq for cell culture batches

```
culture_corrected_data = ComBat_seq(counts = as.matrix(culture_uncorrected[,sample_names]), batch = culture_batch, group = groups)
```

```
## Found 4 batches
## Using full model in ComBat-seq.
## Adjusting for 3 covariate(s) or covariate level(s)
## Estimating dispersions
## Fitting the GLM model
## Shrinkage off - using GLM estimates for parameters
## Adjusting the data
```

Join the gene names onto the now corrected counts from ComBat\_seq

```
culture_corrected_data = cbind(culture_uncorrected[,c("Name")], culture_corrected_data)
```

```
#compare dimensions of corrected and uncorrected data sets
dim(culture_uncorrected)
```

```
## [1] 2632   15
```

```
dim(culture_corrected_data)
```

```
## [1] 2632   15
```

```
#visually compare values of corrected and uncorrected data sets
head(culture_uncorrected)
```

```
head(culture_corrected_data)
```

```
##                    FGF2 FGF3  FGF4 NT1  NT3  NT4 TGFb1 TGFb2 TGFb3 TGFb4 SB1 
## [1,] "let-7a-2-3p" "0"  "0"   "0"  "0"  "0"  "0" "0"   "0"   "0"   "0"   "0" 
## [2,] "let-7a-3p"   "0"  "0"   "0"  "0"  "0"  "0" "0"   "0"   "0"   "0"   "0" 
## [3,] "let-7a-5p"   "1"  "26"  "6"  "85" "72" "9" "13"  "2"   "69"  "24"  "5" 
## [4,] "let-7b-3p"   "0"  "0"   "0"  "0"  "0"  "0" "0"   "0"   "0"   "0"   "0" 
## [5,] "let-7b-5p"   "1"  "102" "3"  "20" "24" "7" "11"  "3"   "29"  "18"  "10"
## [6,] "let-7c-3p"   "0"  "0"   "0"  "0"  "0"  "0" "0"   "0"   "0"   "0"   "0" 
##      SB2 SB3  SB4 
## [1,] "0" "0"  "0" 
## [2,] "0" "0"  "0" 
## [3,] "3" "16" "4" 
## [4,] "0" "0"  "0" 
## [5,] "3" "18" "16"
## [6,] "0" "0"  "0"
```

Convert to dataframe

```
# Convert the character matrix to a data frame
culture_corr_dataframe <- data.frame(culture_corrected_data, stringsAsFactors = FALSE)
```

Write to excel file

```
#library("writexl")
# Set the filename for the Excel file
culture_filename <- paste0(getwd(), "/CultureBatchCorrCounts.xlsx")
# Save the data frame to an Excel file
write_xlsx(culture_corr_dataframe, path = culture_filename)
```
