## Supplemental Code for "Epicardial extracellular vesicles modulate gene expression following ischemia-reperfusion injury in heart-on-a-chip": ComBat_Seq_KW_052923.Rmd

````
---
title: "Batch Effect Adjustment in R Using Combatseq"
output:
  html_document:
    df_print: paged
  pdf_document: default
  word_document: default
---

Citation: Zhang Y, Parmigiani G, Johnson WE. ComBat-seq: Batch effect adjustment for RNA-seq count data. NAR Genomics Bioinforma. 2020;2(3):1-10. doi:10.1093/nargab/lqaa078

Performed By: Karl Wagner

Date: May 29, 2023

Description: Batch effect adjustment of Epi-EV miRNA seq raw counts according to RNA extraction and epicardial culture batches.


# RNA extraction batch effect adjustment
Load necessary packages

```{r}
library("sva")
library("ggplot2")
```
Load the raw miRNA count data 
```{r}
uncorrected_data = read.table("Uncorrected_rawcounts_all4.txt", header=TRUE, sep="\t", as.is=c(1))

#First column - keep as characters
```
```{r}
names(uncorrected_data) = c("Name", "FGF2", "FGF3", "FGF4", "NT1", "NT3", "NT4", "TGFb1", "TGFb2", "TGFb3", "TGFb4", "SB1", "SB2", "SB3", "SB4")
sample_names = names(uncorrected_data)[2:length(names(uncorrected_data))]
```

Review data structure
```{r}
head(uncorrected_data)
dim(uncorrected_data)
```

Define samples, conditions (groups), and batches
```{r}
conditions = c("FGF", "FGF", "FGF", "NT", "NT", "NT", "TGFb", "TGFb", "TGFb", "TGFb", "SB", "SB", "SB", "SB")

#Define RNA extraction batches
RNA_batch = c(1, 2, 2, 1, 2, 2, 1, 1, 2, 2, 1, 1, 2, 2)

#Define epicardial cell culture batches
replicates = c(2, 3, 4, 1, 3, 4, 1, 2, 3, 4, 1, 2, 3, 4)
```

Transform the format of groups and batches from names to numbers
```{r}
#in the command below "sapply" is used to apply the "switch" command to each element and convert names to numbers as we define
#for conditions:
groups = sapply(as.character(conditions), switch, "FGF" = 1, "NT" = 2, "TGFb" = 3, "SB" = 4, USE.NAMES = F)
```
Run Combatseq for RNA extraction batch 
```{r}
corrected_data = ComBat_seq(counts = as.matrix(uncorrected_data[,sample_names]), batch = RNA_batch, group = groups)
```
Join the gene names onto the now corrected counts from ComBat_seq
```{r}
corrected_data = cbind(uncorrected_data[,c("Name")], corrected_data)
```
```{r}
#compare dimensions of corrected and uncorrected data sets
dim(uncorrected_data)
dim(corrected_data)
```
```{r}
#visually compare values of corrected and uncorrected data sets
head(uncorrected_data)
head(corrected_data)
```
Convert to dataframe
```{r}
# Convert the character matrix to a data frame
corr_dataframe <- data.frame(corrected_data, stringsAsFactors = FALSE)
```


Write to excel file

```{r}
library("writexl")
# Set the filename for the Excel file
filename <- paste0(getwd(), "/BatchCorrCounts.xlsx")
# Save the data frame to an Excel file
write_xlsx(corr_dataframe, path = filename)
```

# Cell culture batch effect adjustment
Load the RNA extraction batch effect adjusted count data
```{r}
culture_uncorrected = read.table("RNABatchCorr_Counts_all4.txt", header=TRUE, sep="\t", as.is=c(1))

#First column - keep as characters
```
```{r}
names(culture_uncorrected) = c("Name", "FGF2", "FGF3", "FGF4", "NT1", "NT3", "NT4", "TGFb1", "TGFb2", "TGFb3", "TGFb4", "SB1", "SB2", "SB3", "SB4")
culture_sample_names = names(culture_uncorrected)[2:length(names(culture_uncorrected))]
```

Review data structure
```{r}
head(culture_uncorrected)
dim(culture_uncorrected)
```

Define samples, conditions, and batches as before
```{r}
#same as before - don't need to re-define
#conditions = c("FGF", "FGF", "FGF", "NT", "NT", "NT", "TGFb", "TGFb", "TGFb", "TGFb", "SB", "SB", "SB", "SB")
#RNA_batch = c(1, 2, 2, 1, 2, 2, 1, 1, 2, 2, 1, 1, 2, 2)

#This second step will focus on culture batches as defined below:
culture_batch = c(2, 3, 4, 1, 3, 4, 1, 2, 3, 4, 1, 2, 3, 4)
```
Transform the format of groups and batches from names to numbers
```{r}
#same as before
#in the command below "sapply" is used to apply the "switch" command to each element and convert names to numbers as we define
#for conditions:
#groups = sapply(as.character(conditions), switch, "FGF" = 1, "NT" = 2, "TGFb" = 3, "SB" = 4, USE.NAMES = F)
```

Run Combatseq for cell culture batches
```{r}
culture_corrected_data = ComBat_seq(counts = as.matrix(culture_uncorrected[,sample_names]), batch = culture_batch, group = groups)
```
Join the gene names onto the now corrected counts from ComBat_seq
```{r}
culture_corrected_data = cbind(culture_uncorrected[,c("Name")], culture_corrected_data)
```
```{r}
#compare dimensions of corrected and uncorrected data sets
dim(culture_uncorrected)
dim(culture_corrected_data)
```
```{r}
#visually compare values of corrected and uncorrected data sets
head(culture_uncorrected)
head(culture_corrected_data)
```

Convert to dataframe
```{r}
# Convert the character matrix to a data frame
culture_corr_dataframe <- data.frame(culture_corrected_data, stringsAsFactors = FALSE)
```


Write to excel file

```{r}
#library("writexl")
# Set the filename for the Excel file
culture_filename <- paste0(getwd(), "/CultureBatchCorrCounts.xlsx")
# Save the data frame to an Excel file
write_xlsx(culture_corr_dataframe, path = culture_filename)
```
````
